# Integrative genomics reveals shared and stress-specific adaptive pathways underlying acidic soil-associated metal toxicity in rice

**DOI:** 10.64898/2026.05.31.729167

**Authors:** Sandeep Jaiswal, Anita Kumari, Binay K Singh, Kuldeep Kumar, Saurav Kumar, Santosh Kumar, Simardeep Kaur, Nitish Ranjan Prakash, Pankaj Baiswar, Alka Bharati, Manjeet Talukdar, Sanjay Behera

## Abstract

Soil acidity-associated toxicities of aluminum (Al), cadmium (Cd), and manganese (Mn) severely constrain rice productivity in upland ecosystems. To investigate the genomic basis of adaptation to acidic soil-related metal stress, we conducted an integrated meta-QTL (M-QTL) and functional genomics analysis in rice. Meta-analysis of 681 QTLs and MTAs from 53 QTL mapping and GWAS studies identified 79 robust M-QTLs, including ten overlapping regions associated with Al-, Cd-, and Mn-responsive traits. A multi-criteria prioritization framework identified 98 candidate genes supported by positional overlap, transcriptomic recurrence, and functional annotation, enriched for ion transport, detoxification, and redox regulation pathways. M-QTL10.9 emerged as a major hotspot enriched for glutathione-S-transferase genes, whereas M-QTL9.5 contained the highest density of prioritized candidates linked to Al and Cd responses. Comparative physiological & biochemical analyses of the contrasting rice genotypes Sahasarang and IR64 revealed genotype-dependent differences in antioxidant responses, metal partitioning, metabolic regulation, and cell wall remodeling under individual and combined metal stresses. Expression profiling of prioritized candidate genes, including OsACO family genes, *OsZIP10*, and *OsGSTU10*, further revealed genotype-dependent transcriptional divergence under combined stress. The identification of overlapping M-QTLs across Al, Cd, and Mn datasets suggests both shared and stress-specific adaptive responses to acidic soil-associated metal stress in rice.

## 1. Introduction

Rice (*Oryza sativa* L.) is a staple food for the majority of the global population, contributing over 20% of daily caloric intake worldwide (Chu & Yu, 2020). Its production must increase by approximately 40% by 2050 to meet future food demands (Milovanovic & Smutka, 2017). However, rice cultivation in upland and marginal environments is increasingly constrained by soil acidity and associated metal toxicities. Approximately one-third of global rice-growing areas, including irrigated (25.6 mha), rainfed lowlands (18.5 mha), and uplands (7.5 mha), are affected by poor soil quality, of which at least 27.1 mha with soil pH below 5.5 is impacted by acidity-related constraints (Haefele *et al*., 2014). In acidic soils, aluminum (Al) and manganese (Mn) toxicity frequently co-occur with phosphorus (P) deficiency (Zhao *et al*., 2014). This often drives phosphatic fertilizer application, nearly 85% of which is derived from sedimentary phosphate rock containing up to 150 mg Cd kgL¹, thereby contributing to cadmium (Cd) accumulation in agricultural soils (Bailey *et al*., 1995; Jung, 2008; Sarwar *et al*., 2010). Soil acidity further increases Cd bioavailability, a process exacerbated by wastewater irrigation and animal manure application (Gao *et al*., 2022). These constraints are expected to intensify under climate change, as altered precipitation regimes are likely to enhance soil acidification and metal mobility in rice-growing environments (Nel *et al*., 2022; Bibi & Rehman, 2023; Corami, 2023; Raza *et al*., 2025). In acidic soils, toxic concentrations of Al, Mn, and Cd severely impair rice growth through distinct but partially overlapping morphological, physiological, and cellular alterations. Al toxicity predominantly suppresses cell division and elongation in the root apex, leading to root thickening and darkening, reduced branching and root hair formation, and, under prolonged exposure, cracking of root apices (Sivaguru *et al*., 1998; Vitorello *et al*., 2005; Kochian *et al*., 2015; Miyasaka *et al*., 2007; Wang *et al*., 2023a). Mn toxicity, which frequently co-occurs with Al stress, primarily affects shoot growth and function by impairing root and shoot elongation, inducing chlorotic and necrotic lesions, disrupting photosynthesis, and reducing productivity (Nelson, 1983; Page *et al*., 2006; Arya and Roy, 2011; Weng *et al*., 2013; Rojas-Lillo *et al*., 2014). In contrast, Cd predominantly accumulates in roots, where it disrupts osmotic regulation, reduces stomatal conductance and transpiration, and causes tissue necrosis and cellular degradation, ultimately resulting in chlorosis, growth inhibition, and impaired plant development (Hermans *et al*., 2011; Gallego *et al*., 2012; Rizwan *et al*., 2016; Abbas *et al*., 2017; Chellaiah, 2018; Kubier *et al*., 2019). At the cellular level, Al, Mn, and Cd primarily disrupt membranes and metabolism via oxidative stress. Al^3^□ binds plasma membrane phospholipids, increasing rigidity, while concurrent Ca deficiency destabilizes cell walls and nucleic acids, impairing water and nutrient uptake and heightening stress sensitivity (Čiamporová, 2002; Jones *et al*., 2006; Inostroza-Blancheteau *et al*., 2008). Mn similarly induces oxidative degradation of biomolecules, whereas Cd-generated reactive oxygen species (ROS) damage membranes and organelles, collectively compromising cellular metabolism and potentially leading to cell death (St. Clair and Lynch, 2005; Fernando *et al*., 2013; Abbas *et al*., 2017). Interactions among Al, Mn, and Cd further modulate toxicity. Cd alters Al partitioning by maintaining root accumulation but reducing shoot translocation, whereas Al can antagonistically suppress Mn uptake (Rees and Sidrak, 1961; Clark, 1977; Blair and Taylor, 1997; Taylor *et al*., 1998; Guo *et al*., 2004; Yang *et al*., 2009; Wang *et al*., 2015). Collectively, toxic levels of these metals disrupt the availability and uptake of essential micro- and macronutrients, leading to chlorosis and morphological abnormalities. Both synergistic and antagonistic metal-nutrient interactions have been reported, with Cd notably interfering with Mn, Ca, P, Mg, K, and N uptake (Moroni *et al*., 2003; Rosas *et al*., 2007; Mora *et al*., 2009; Führs *et al*., 2010; Nazar *et al*., 2012; Xu *et al*., 2017). Recent reviews comprehensively summarize the physiological and molecular responses to these toxicities (Rasheed *et al*., 2021; Tang *et al*., 2023; Wang *et al*., 2023b; Haider *et al*., 2023). These synergistic and antagonistic interactions indicate that metal stress responses involve overlapping regulatory, detoxification, and ion homeostasis pathways despite substantial differences in downstream physiological outcomes. Although Al, Cd, and Mn toxicities frequently co-occur in acidic soils, their primary toxicodynamic targets differ substantially. Aluminum toxicity predominantly affects root apical growth and cell wall integrity, whereas Cd and Mn stresses are more strongly associated with metal uptake, translocation, oxidative imbalance, and shoot-associated injury. Consequently, genomic regions associated with these stresses may reflect distinct but interconnected adaptive processes operating across different physiological contexts. Nevertheless, overlap in downstream detoxification, ion homeostasis, oxidative buffering, and stress-signaling pathways may contribute to broader adaptation under acidic soil conditions.

Numerous QTLs associated with root growth, metal uptake, translocation, and stress adaptation under Al, Cd, and Mn toxicity have previously been identified in rice using both biparental and natural populations. For Al tolerance, QTLs have been reported across all 12 chromosomes, several of which explain substantial phenotypic variation within individual populations (Nguyen *et al*., 2001, 2003; Ma *et al*., 2002; Xue *et al*., 2006, 2007; Wu *et al*., 2000). Similarly, major loci associated with Cd tolerance and accumulation have been identified, including large-effect QTLs controlling shoot and grain Cd accumulation on chromosome 11, together with additional loci on chromosomes 3 and 7 associated with metal uptake, partitioning, and stress adaptation (Koike *et al*., 2004; Nakanishi *et al*., 2006; Uraguchi *et al*., 2011; Ishikawa *et al*., 2012; Luo *et al*., 2018; Yan *et al*., 2019). In contrast, Mn toxicity tolerance and accumulation have been comparatively less explored (Wang *et al*., 2002; Liu *et al*., 2017; Shrestha *et al*., 2018). However, substantial heterogeneity exists among these studies with respect to trait definitions, developmental stages, stress exposure systems, mapping populations, and phenotyping strategies, thereby complicating direct biological interpretation across datasets. Differences in experimental design, environments, parental backgrounds, population sizes, marker density, and statistical methodologies have further affected QTL consistency and associated phenotypic variance, while fine mapping of most causal genes remains incomplete. Consequently, the identification of stable and recurrent genomic regions associated with diverse but interconnected metal stress responses remains essential for understanding adaptation to acidic soil environments and for improving marker-assisted breeding strategies. Meta-QTL (M-QTL) analysis provides an effective framework to integrate independent studies, refine confidence intervals, and identify robust markers and candidate genes associated with stress adaptation in crops (Goffinet & Gerber, 2000; Kumari *et al*., 2023 & 2024). In this study, we hypothesized that stable genomic regions associated with interconnected adaptive responses to Al, Cd, and Mn toxicity can be identified through integration of independent genetic studies. Accordingly, we implemented an integrative meta-QTL framework to project published QTLs and MTAs onto a consensus genetic map, define robust M-QTL intervals, and prioritize candidate genes through downstream in silico analyses. Comparative evaluation of the contrasting rice genotypes Sahasarang and IR64 under individual and combined metal stresses, together with expression, morphophysiological, biochemical, ionomic, and cell wall compositional analyses, was performed to characterize genotype-dependent responses to complex metal stress and to examine the expression behaviour of prioritized candidate genes. Collectively, these findings provide biologically prioritized targets and a physiological framework for future functional studies. Collectively, these findings provide biologically prioritized targets and mechanistic insights for functional genomics and breeding of rice adapted to acidic soil-associated metal stress environments.

## 2. Materials and methods

### 2.1. Methodology overview

A schematic overview of the experimental design and analytical workflow employed in the present study is provided in **Fig. 1**.

**Fig. 1.**
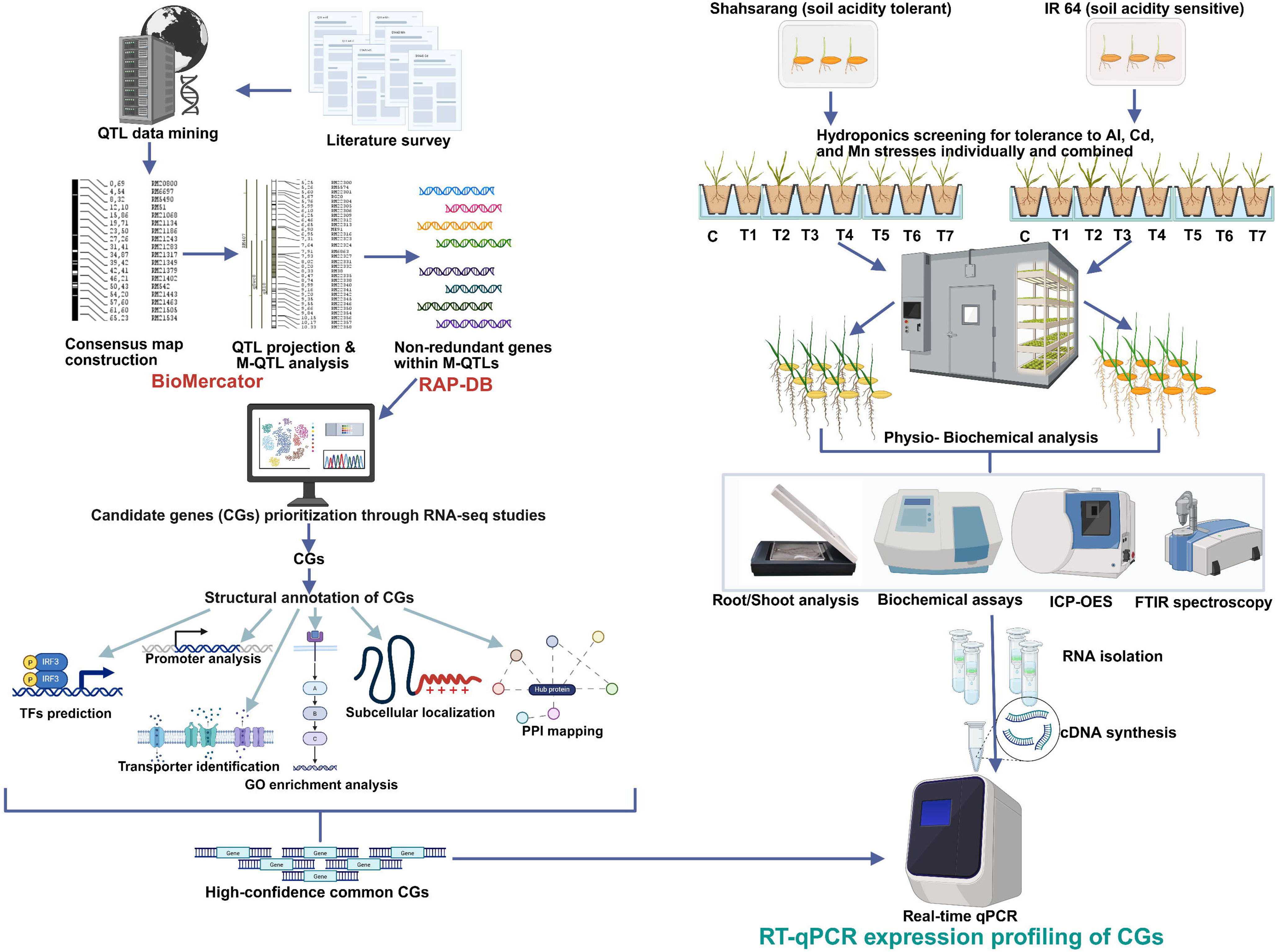
Schematic overview of the methodology employed for the study.

### 2.2. Literature survey, QTL data mining, and preparation of input files for M-QTL analysis

A comprehensive literature survey was conducted to compile QTL mapping and genome-wide association studies (GWAS) associated with Al, Cd, and Mn toxicity tolerance in rice published up to December 2024. Information including QTL identifiers, associated traits, chromosomal positions, logarithm of odds (LOD) scores, phenotypic variance explained (R²), genetic positions (cM), and confidence intervals (CI) was retrieved from the selected studies. When LOD scores or R² values were unavailable, default values of 3.0 and 10%, respectively, were assigned following established guidelines (Khahani *et al*., 2019; Jaiswal *et al*., 2026).. For GWAS datasets, physical marker positions were converted to centiMorgan (cM) positions using a conversion factor of 200 kb cML¹ based on the average recombination rate of the rice genome (Orjuela *et al*., 2010). SNP markers were subsequently aligned to the rice reference linkage map developed by Temnykh *et al*. (2001) by assigning them to the nearest flanking markers. A symmetrical confidence interval of ±2 cM was applied to these loci to define QTL boundaries for downstream meta-analysis (Jaiswal *et al*., 2024; Jaiswal *et al*., 2026). Genetic maps were reconstructed for individual studies by organizing marker positions within linkage groups to generate standardized input files for M-QTL analysis.

### 2.3. Consensus genetic map construction and QTL meta-analysis

A consensus genetic map was constructed by integrating genetic map information from all selected studies with the rice reference linkage map using BioMercator v4.2.3 (Temnykh *et al*., 2001; Arcade *et al*., 2004; Sosnowski *et al*., 2012). The resulting consensus map served as the reference framework for projecting all shortlisted QTLs and marker–trait associations (MTAs). To facilitate identification of recurrent genomic regions associated with interconnected manifestations of acidic soil-associated metal stress adaptation, all projected loci were integrated under a unified trait framework designated as “AlCdMn stress-associated traits,” while retaining their original trait annotations for downstream biological interpretation. Meta-QTL analysis was subsequently performed using the Veyrieras two-step algorithm implemented in BioMercator (Veyrieras *et al*., 2007; Kumari *et al*., 2023).

### 2.4. Candidate gene prioritization across M-QTL intervals

Physical coordinates of markers flanking the projected M-QTLs were used to retrieve all genes located within the 79 M-QTL intervals from the Rice Annotation Project Database (RAP-DB) (Sakai *et al*., 2013). The resulting non-redundant gene set was subjected to comparative transcriptomic analysis by cross-referencing with fourteen publicly available root-specific RNA-seq datasets describing rice responses to Al, Cd, and Mn stresses (Awasthi *et al*., 2021; Arenhart *et al*., 2014; Arbelaez *et al*., 2017; Gallo-Franco *et a*l., 2023; Cao *et al*., 2019; Fan *et al*., 2021; Li *et al*., 2017; Liu *et al*., 2020; Sun *et al*., 2019; Wang *et al*., 2021; Wang *et al*., 2022; Yu *et al*., 2021; He & Li, 2022; Dong *et al*., 2024) using reference (RGAP, version 7 - https://rice.uga.edu/download_osa1r7.shtml) based assembly approach following Kumar *et al*., (2021). The resulting raw count matrix was used to estimate gene-level statistics, including logL fold change (logLFC), standard error of the logL fold change (lfcSE), and associated P-values using the DESeq2 package in R. Genes lacking valid logL fold change or standard error estimates were excluded from downstream meta-analysis to ensure reliable variance estimation. No pre-filtering based on statistical significance was applied at this stage to minimize selection bias across datasets. All processed study-level datasets were consolidated into a single expression matrix and subjected to random-effects meta-analysis to account for both within-study variance and between-study heterogeneity. Meta-analysis was performed independently for each gene using the MetavolcanoR package implemented in R. For each gene, logL fold change estimates from all available studies were integrated using a random-effects model, thereby accommodating variability arising from differences in experimental design, stress conditions, and transcriptomic platforms across independent datasets. To refine biologically relevant candidate genes, a quantitative weightage-based prioritization framework was implemented. Each gene was assigned a composite prioritization score (CPS) integrating transcriptomic recurrence (E), multimetal responsiveness (F), functional relevance (G), and M-QTL positional strength (H) according to the following equation:

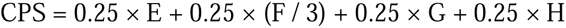

where E represents the number of independent transcriptomic datasets reporting differential expression, F denotes responsiveness across Al, Cd, and Mn stress conditions, G reflects curated functional relevance based on annotation and published evidence, and H represents M-QTL positional strength calculated as: H = 0.333 × R² + 0.333 × C + 0.333 × T

where R² corresponds to normalized phenotypic variance explained, C represents QTL cluster size, and T denotes trait breadth within each M-QTL interval. Equal weighting was applied to balance expression evidence, stress breadth, biological plausibility, and genetic robustness while minimizing overrepresentation of any single evidence layer. Genes exceeding the prioritization threshold were retained as high-confidence candidates for downstream *in silico* characterization and expression profiling. Gene ontology enrichment analysis of prioritized CGs was performed using ShinyGO v0.80 (Ge *et al*., 2020).

### 2.5. Investigation of the prioritized CGs2.5.1. Protein–protein interaction (PPI) mapping

To investigate functional interactions among the prioritized candidate genes (CGs), protein–protein interaction (PPI) analysis was performed using the STRING database (https://string-db.org) with Oryza sativa selected as the reference organism. Candidate genes were queried using a minimum interaction confidence score of ≥0.4, and the resulting interaction network was visualized in Cytoscape v3.9.1. Hub genes were identified using the CytoHubba plugin based on Maximal Clique Centrality (MCC) scores and overall network connectivity. Functional enrichment analysis, including Gene Ontology (GO) and Kyoto Encyclopedia of Genes and Genomes (KEGG) pathway analyses, was subsequently performed to identify biologically enriched pathways and functionally interconnected stress-responsive modules associated with Al, Cd, and Mn stress adaptation.

#### 2.5.2. Promoter analysis

To investigate potential cis-regulatory signatures associated with Al, Cd, and Mn stress responses, in silico promoter analysis of CGs within the identified M-QTL intervals was performed using 2 kb upstream sequences from the translational start codon (ATG). These sequences were retrieved from the Rice Annotation Project Database (RAP-DB; https://rapdb.dna.affrc.go.jp). Cis-regulatory element analysis was performed using the PlantCARE database (Lescot *et al*., 2002). All detected cis-elements were grouped into functional classes to reduce redundancy and facilitate interpretation. These included ABA-responsive (ABRE, ABRE2, ABRE3a, ABRE4), MYB-binding (MYB, MBS, MYB recognition sites), MYC/bHLH-associated (MYC/Myc), dehydration-responsive (DRE, DRE core, DRE1), jasmonate/ethylene-responsive (TGACG- and CGTCA-motifs), auxin-responsive (AuxRR-core and TGA-element), light-responsive (G-box, ACE, Box 4, I-box), defense-related (W-box and WRE3), and core promoter elements (TATA- and CAAT-boxes). Transcript isoforms were collapsed to the gene level, and motif counts were summed to generate gene-wise cis-element profiles. A gene × motif-class matrix was constructed and visualized using heatmaps to compare promoter architectures among CGs.

#### 2.5.3. Identification of transcription factors-, membrane transporter-encoding CGs and prediction of their subcellular localization

Candidate genes encoding TFs among the selected CGs were identified using the PlantTFDB v5.0 web server (http://planttfdb.gao-lab.org/prediction.php). Additionally, subcellular localization was predicted using DeepLoc v2.1 (Thumuluri *et al*., 2022), and transmembrane domains were identified using DeepTMHMM v1.0 (Hallgren *et al*., 2022).

### 2.6. Evaluation of contrasting rice genotypes Sahasarang and IR64 under Al, Cd, and Mn stress conditions

Sahasarang and IR64 were selected as contrasting rice genotypes exhibiting differential responses to acidic soil-associated metal stresses in previous studies. Sahasarang is a high-yielding variety that has frequently been reported as relatively adapted to acidic soil environments and Al-dominant stress conditions and has been used as a tolerant reference genotype in germplasm screening programs targeting acidic soil adaptation (Pal *et al*., 2011; Debnath *et al*., 2017; Devi *et al*., 2018; Raj *et al*., 2025; Sonu *et al*., 2025; Balakrishnan *et al*., 2025). In contrast, IR64 is widely used as a stress-sensitive reference genotype in studies involving Al, Cd, Mn, and Fe toxicity (Ma *et al*., 2007; Dubey *et al*., 2014; Pandey and Gupta, 2018; Raghuvanshi *et al*., 2022; Maity *et al*., 2025).

#### 2.6.1. Imposition of Al, Cd, and Mn stress and sample collection

Sterilized seeds of Sahasarang and IR64 were germinated on autoclaved, water-moistened Whatman No. 1 filter paper at 25°C in darkness for 96 h. Seedlings were subsequently transferred to a hydroponic floating raft system containing modified Magnavaca nutrient solution (Jaiswal *et al.,* 2023) and maintained in a Conviron growth chamber at 25 ± 1°C under a 16 h light/8 h dark photoperiod with a light intensity of 250 µmol mL² sL¹ for 10 days. Metal stress treatments consisted of 200 µM AlClL, 30 µM CdClL, and 300 µM MnClL applied individually and in combination. Eight treatment combinations (control, Al, Cd, Mn, Al+Cd, Al+Mn, Cd+Mn, and Al+Cd+Mn) were evaluated per genotype using three independent replicate trays per treatment. After 10 days of stress exposure, three randomly selected seedlings from each replicate were harvested, washed with autoclaved double-distilled water, and processed for subsequent analyses.

#### 2.6.2. Quantitative analysis of root morphology and biomass

Cleaned root samples were scanned at high resolution using an Epson Perfection V-700 flatbed scanner and analyzed with WinRHIZO Pro software (Regent Instruments Inc., Canada) to quantify root morphological traits, including total root length (RL), root surface area (SA), number of forks (FA), root volume (RV), and average root diameter (Avg_D). Fresh root and shoot biomass were recorded immediately after harvest using an analytical balance to obtain root fresh weight (FW_R) and shoot fresh weight (FW_S). Samples were then oven-dried at 70°C for 72 h to determine root dry weight (DW_R) and shoot dry weight (DW_S).

#### 2.6.3. Assessment of physiological and biochemical responses of root and shoot

Total phenolic content (TPC) was quantified using the Folin–Ciocalteu colorimetric method (Singleton & Rossi, 1965), while antioxidant capacity was evaluated using the ferric reducing antioxidant power (FRAP) assay (Benzie & Strain, 1996). Total soluble sugars (TSS) were estimated using the anthrone method (Yemm & Willis, 1954), and starch content was determined following enzymatic hydrolysis and colorimetric measurement (McCready *et al*., 1950).

#### 2.6.4. Cell wall compositional profiling and metal accumulation analysis in root and shoot tissues

Cell wall composition of root and shoot tissues was analyzed by Fourier-transform infrared spectroscopy (FTIR) using the KBr pellet method (Alonso-Simón *et al*., 2011; Largo-Gosens *et al*., 2014). Oven-dried samples (70°C, 72 h) were finely ground, mixed with spectroscopic-grade KBr (1:100, w/w), and pressed into pellets. Spectra were recorded on a Bruker VERTEX 70 spectrometer over 4000–400 cmL¹ at 4 cmL¹ resolution with 32 scans. Baseline correction and noise reduction were performed using OMNIC 32, and spectra were processed in Origin 8.6 to evaluate characteristic cell wall components.

For metal accumulation analysis, 100 mg of the same oven-dried, ground root and shoot samples were subjected to microwave-assisted digestion with a 2:1 (v/v) HNOL:HClOL mixture (Zarcinas *et al*., 1987). Digests were filtered (Whatman No. 42) and diluted to 50 ml with ultrapure water (dilution factor = 1000). Al, Cd, and Mn concentrations were quantified using ICP-OES (PerkinElmer Avio 500) and expressed on a dry weight basis for genotype- and treatment-wise comparisons.

### 2.7. RT-qPCR-based expression profiling of CGs

RT-qPCR analysis was performed in Shahsarang and IR64 using root and leaf tissues from control and metal-treated seedlings, following the methodology described in Jaiswal *et al*. (2024). Briefly, total RNA was isolated, DNase-treated, and reverse-transcribed to cDNA as previously described. Gene-specific primers were used to quantify transcript levels of prioritized CGs on a CFX96 Real-Time PCR System (Bio-Rad, USA) using TB Green® Premix Ex Taq™ II (Takara Bio). Each reaction was run in three technical replicates, with rice β-actin serving as the internal reference gene. Relative expression levels were calculated using the 2^–ΔΔCt method (Schmittgen and Livak, 2008). Primer sequences are provided in **Table S1**.

### 2.8. Statistical analysis

All data are presented as mean ± SE. Treatment effects were assessed by one-way ANOVA, followed by Tukey’s HSD test (P < 0.05) for mean separation, with significant differences denoted by letter groupings (Gomez and Gomez, 1984). Correlation analysis was used to examine pairwise trait relationships. Principal component analysis (PCA) was conducted on z-score–standardized root–shoot morphological and biomass traits; components with eigenvalues > 1 were retained, and loadings and biplots were used to identify genotype- and tissue-discriminating traits. Statistical analyses were performed in R (v4.3.0) using the stats and agricolae packages, while data preprocessing and visualization were carried out in Python (v3.11) using pandas, numpy, and matplotlib.

## 3. Results

### 3.1. Survey of QTL and GWAS studies associated with Al, Cd, and Mn stress responses in rice

A systematic literature survey conducted up to December 2024 identified 53 studies, including 37 biparental QTL mapping studies and 16 GWAS, investigating the genetic architecture of Al-, Cd-, and Mn-associated stress responses in rice. A total of 681 QTLs and marker–trait associations (MTAs) were compiled together with their chromosomal positions, confidence intervals (CI), logarithm of odds (LOD) scores, and phenotypic variance explained (PVE). The integrated dataset encompassed substantial variation in stress phenotypes, developmental stages, tissues, population structures, and experimental systems, thereby capturing multiple dimensions of metal stress adaptation in rice. Phenotypic evaluations differed according to the dominant physiological effects of each metal stress. Al-associated studies primarily focused on root growth–related traits, including relative root elongation (RRE), relative root length (RRL), and root biomass, reflecting the rapid inhibition of root apical growth under Al toxicity. In contrast, Cd- and Mn-associated studies more frequently evaluated tissue-specific metal accumulation, translocation efficiency, grain metal concentration, and shoot-associated toxicity responses. Several studies additionally incorporated biomass, nutrient homeostasis, and physiological traits to capture broader stress-associated responses (**Table S2**). Despite this phenotypic diversity, many measured traits converged on common adaptive processes associated with ion homeostasis, metal sequestration, detoxification efficiency, and maintenance of root growth under stress conditions. The retrieved studies exhibited substantial methodological variation, including differences in molecular marker systems, population structures, mapping resolution, and environmental conditions. Biparental populations ranged from 39 to 373 individuals, whereas GWAS panels included 127–873 accessions **(Table S2)**. Consequently, the number of reported loci per study varied widely (1–119), with confidence intervals ranging from 0.01 to 77.45 cM and PVE ranging from 0.11% to 92%. The identified loci were distributed across all 12 rice chromosomes, although their density varied considerably, with chromosome 3 harboring the highest number of loci and chromosome 10 the lowest **(Fig. S1)**. The broad heterogeneity and partial overlap among studies highlighted the need for consensus mapping and meta-QTL analysis to refine genomic intervals and identify robust loci associated with coordinated adaptation to multi-metal stress conditions.

### 3.2. Consensus map construction and meta-analysis of metal stress-associated QTLs

A consensus genetic map comprising 20,001 markers across 1837.97 cM was constructed by integrating data from 53 QTL mapping and GWAS studies. Of the 681 retrieved QTLs and MTAs, 493 (72.39%) were successfully projected onto the consensus map, indicating substantial marker correspondence among independent studies. Meta-analysis was subsequently performed within a unified genomic framework while retaining the original trait annotations to preserve biological context during downstream interpretation. Meta-analysis identified 83 M-QTLs distributed across all 12 rice chromosomes, of which 79 contained ≥2 overlapping loci and collectively represented 306 projected QTLs (**Table S3**). The number of retained M-QTLs ranged from three on chromosome 2 to ten on chromosomes 7 and 10. Notably, ten M-QTLs (1.7, 3.5, 6.2, 6.7, 6.8, 7.1, 7.3, 10.8, 10.9, and 11.7) were associated with Al-, Cd-, and Mn-responsive traits, identifying recurrent genomic regions potentially linked to partially convergent adaptive responses across multiple metal stress conditions. Given the diversity of underlying phenotypes and experimental systems, these overlapping loci were interpreted as shared stress-associated genomic regions rather than direct evidence of identical tolerance mechanisms. Among the retained M-QTLs, 23 harboured ≥5 overlapping loci, whereas five regions (6.4, 7.2, 7.3, 7.4, and 8.1) contained ≥8 projected loci, with a maximum cluster size of 12. Several regions, including M-QTLs 7.2, 7.4, 9.2, 9.3, 9.7, and 11.7, were additionally associated with relatively high phenotypic variance explained (14.80–38.25%), supporting their stable recurrence across independent studies. Meta-analysis substantially improved mapping resolution, reducing the average confidence interval from 6.90 cM to 1.85 cM and facilitating downstream prioritization of stress-responsive candidate genes. The ten multimetal-associated M-QTLs exhibited an average cluster size of 5.6 and a mean confidence interval of 1.59 cM (**Table S4**). Collectively, the identified regions revealed substantial convergence among independently reported metal-responsive loci despite pronounced phenotypic and methodological variation across studies, highlighting several recurrent genomic intervals potentially associated with coordinated adaptation to acidic soil-associated metal stress.

### 3.3. Candidate gene prioritization within M-QTL regions

A total of 4,702 non-redundant genes were identified within the physical intervals of the 79 retained M-QTLs (**Table S5**). Integration with fourteen publicly available root-specific RNA-seq datasets associated with Al, Cd, and Mn stress responses identified 705 genes exhibiting differential expression under at least one metal stress condition (**Fig. 2b**). To refine biologically relevant stress-associated loci, positional overlap, stress-responsive transcriptional behaviour, and functional annotations were subsequently integrated through a composite prioritization framework, resulting in the identification of 98 high-confidence CGs with prioritization scores ranging from 0.45 to 0.87 (**Table S6**).

**Fig. 2.**
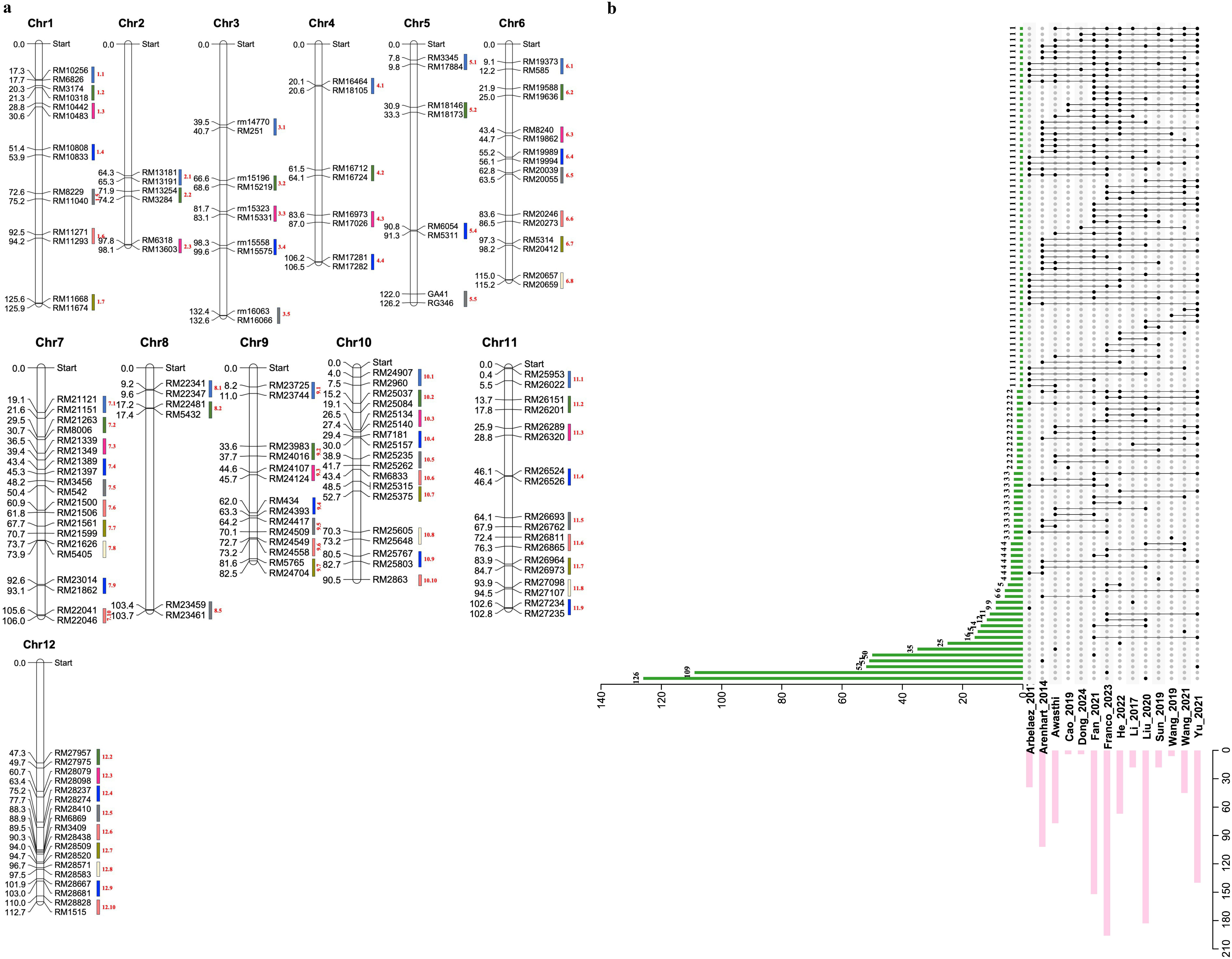
**a**. Distribution of 79 meta-QTLs (M-QTLs), each supported by at least two overlapping QTLs associated with AlCdMn toxicity tolerance, across the 12 rice chromosomes. **b**: An upset plot highlighting the number of differentially regulated genes (705) identified in 14 rice Al-Cd-Mn-toxicity tolerance-related transcriptomic studies.

The prioritized candidate set showed strong functional enrichment for genes associated with membrane transport, redox homeostasis, hormonal signalling, cellular detoxification, and stress-responsive regulatory pathways. Several prioritized loci encoded transporters, glutathione S-transferases, peroxidases, transcription factors, and signalling-associated proteins implicated in ion homeostasis, oxidative stress buffering, and adaptive stress responses.

### 3.4. In silico functional characterization of prioritized candidate genes

#### 3.4.1. Functional annotation and enrichment of prioritized CGs

Functional annotation and GO enrichment analysis of the 98 prioritized candidate genes (CGs) revealed significant overrepresentation of pathways associated with redox regulation, glutathione-dependent detoxification, oxidoreductase activity, and metal stress-responsive cellular processes (FDR < 0.05). Enriched biological processes included oxidative stress response, glutathione metabolism, hydrogen peroxide catabolism, and chemical detoxification, highlighting the strong representation of antioxidant and stress-buffering functions within recurrent M-QTL intervals (**Fig. S2**).

Independent keyword enrichment analysis further supported the predominance of oxidative stress-, detoxification-, and metal ion-associated functional categories. Collectively, these analyses identified recurrent enrichment of redox homeostasis and cellular detoxification pathways across prioritized genomic regions, supporting their potential involvement in coordinated adaptation to acidic soil-associated metal stress conditions.

#### 3.4.2. Protein–protein interaction (PPI) network analysis

Protein–protein interaction analysis identified a highly interconnected network enriched for pathways related to redox regulation, detoxification, metabolic adaptation, and protein quality control (**Fig. 3**; **Table S7**). CytoHubba analysis identified ten hub genes, among which *Os09g0440300*, *Os09g0453800*, *Os10g0390500*, *Os07g0638400*, and *Os11g0484000* exhibited the highest maximal clique centrality (MCC) scores, indicating prominent positions within the interaction network.

**Fig. 3.**
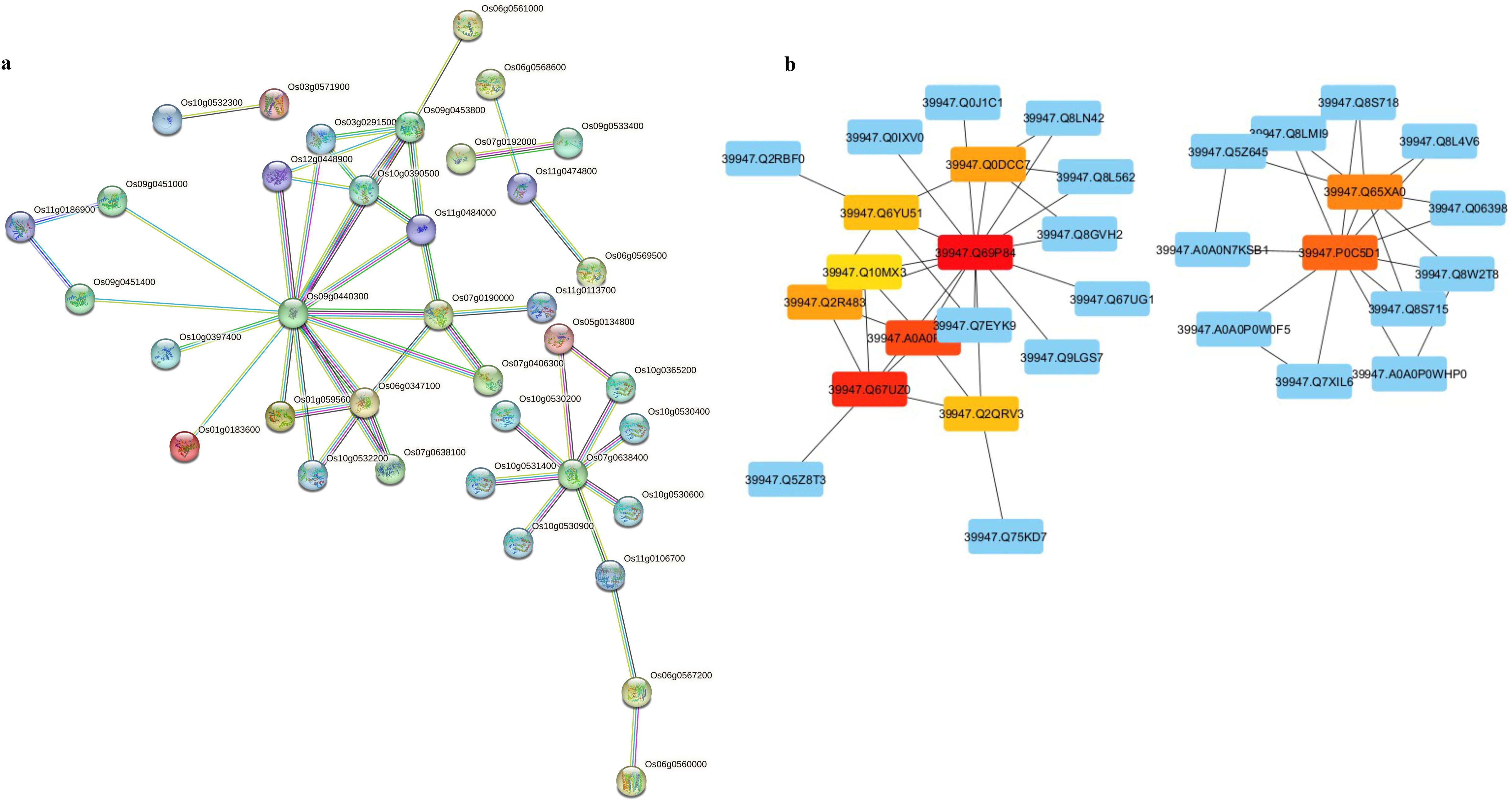
**a.** Protein–protein interaction network of 98 prioritized CGs responsive to multimetal stress in rice. **b.** CytoHubba analysis of the STRING-derived PPI network identified ten hubs.

The interaction architecture revealed strongly interconnected functional modules centered on antioxidant defense, glutathione-dependent detoxification, and maintenance of cellular homeostasis under metal stress conditions. Subnetworks enriched for 1-Cys peroxiredoxins, glutathione S-transferases, and aldehyde dehydrogenases formed major interaction clusters linked to ROS buffering and detoxification processes. Additional modules included proteins associated with membrane stability, metabolic adjustment, isoprenoid biosynthesis, and ubiquitin-mediated protein turnover, supporting the coordinated involvement of redox regulation, metabolic plasticity, and protein quality control pathways in adaptation to acidic soil-associated metal stress.

#### 3.4.3. Promoter cis-regulatory element analysis of prioritized candidate genes

Promoter regions (2 kb upstream of the transcription start site) of the 98 prioritized candidate genes (CGs) were analyzed to examine the cis-regulatory architecture associated with acidic soil-related metal stress responses. Cis-element analysis revealed widespread enrichment of stress-, hormone-, and transcription factor-associated motifs across prioritized CG promoters (**Fig. 4; Table S8**). Among the detected motifs, MYB- and MYC-recognition elements were the most abundant and broadly distributed, followed by ABA-responsive elements (ABREs), dehydration-responsive elements (DREs), and jasmonate/ethylene-associated motifs. W-box elements associated with WRKY transcription factor binding were identified in a smaller subset of stress-responsive genes. The recurrent occurrence of these motifs across prioritized loci supports the potential involvement of ABA-, ROS-, and stress-associated transcriptional regulatory pathways under metal stress conditions.

**Fig. 4.**
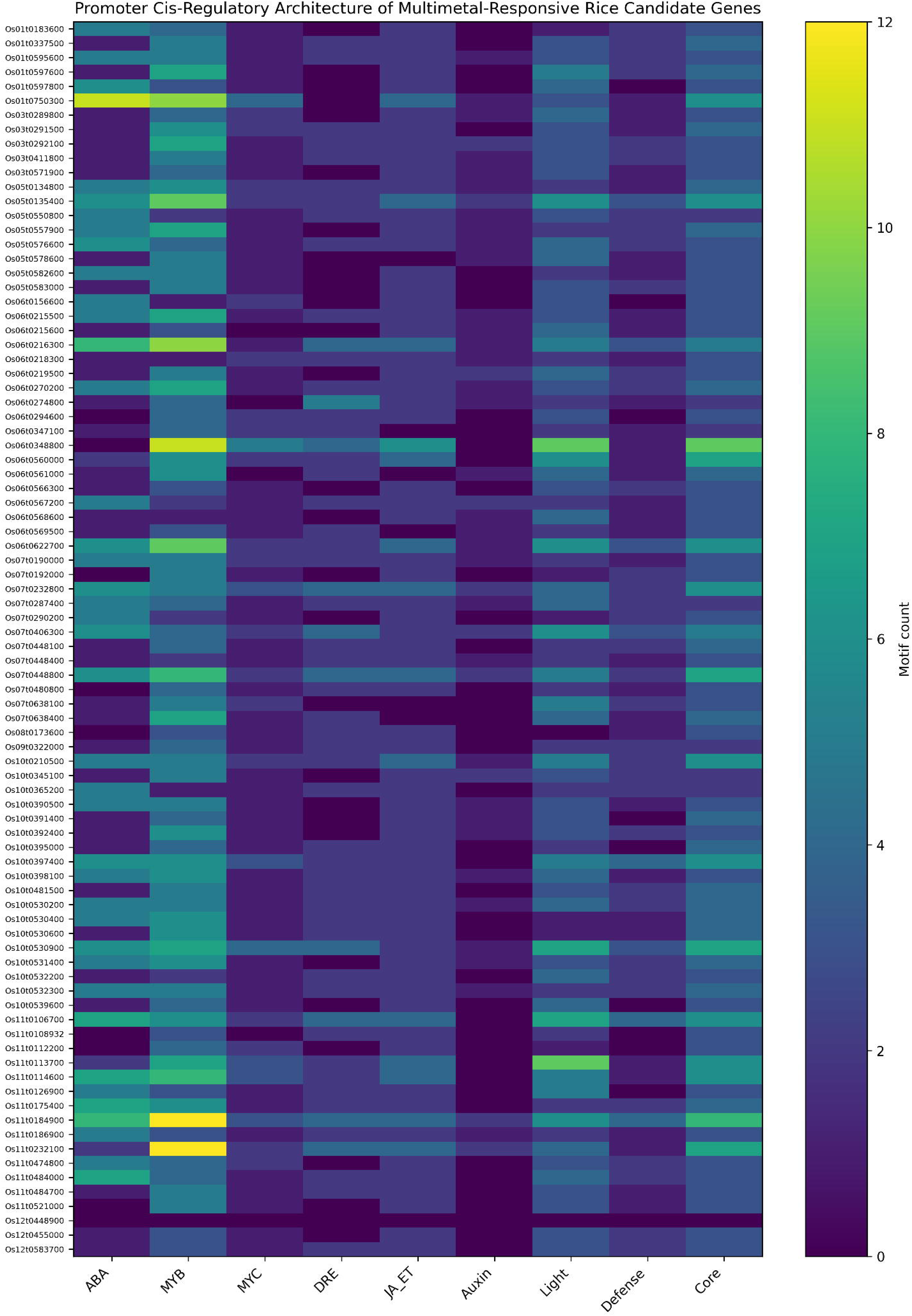
*In silico* identification of regulatory elements within the putative promoter regions of the identified 98 prioritized candidate genes. Heatmap depicts the distribution of major functional cis-regulatory motif classes within the 2-kb upstream promoter regions. Cis-elements were grouped into hormone-responsive (ABA, JA/ET, auxin), transcription factor binding (MYB, MYC), stress-responsive (DRE), light-responsive, defense-associated, and core promoter categories. Color intensity represents absolute motif counts per gene.

Several candidate genes, including *Os11g0184900*, *Os11g0232100*, *Os06g0216300*, and *Os01g0750300*, exhibited relatively high densities of MYB-, ABA-, and jasmonate-responsive motifs, frequently accompanied by ethylene-associated regulatory elements. In contrast, transporter-associated genes such as OsZIP10, OsCOPT7, and OsHKT4 displayed comparatively enriched stress-and ion-responsive *cis*-elements (**Table S8**). Overall, the promoter architectures of prioritized CGs revealed extensive representation of stress- and hormone-responsive regulatory motifs within recurrent M-QTL intervals, supporting the possibility of coordinated transcriptional regulation during adaptation to acidic soil-associated metal stress.

#### 3.4.4. Transcriptional regulation, membrane transport, and compartmentalized stress adaptation within recurrent M-QTL regions

Functional annotation of prioritized candidate genes (CGs) revealed extensive enrichment of transcriptional regulators, membrane-associated transport proteins, and compartment-specific adaptive components within recurrent M-QTL intervals (**Tables S9–S11**). These functional categories collectively support coordinated regulation of ion homeostasis, ROS buffering, cellular signaling, and structural adaptation under acidic soil-associated metal stress conditions.

Multiple prioritized CGs encoded transcription factors belonging to NAC, AP2/ERF, bHLH, WRKY, bZIP, and G2-like families, indicating substantial stress-responsive transcriptional reprogramming capacity within recurrent genomic intervals. Representative regulators included *OsNAC5*, *OsERF102*, *OsbHLH120*, *OsWRKY8*, and *OsbZIP60*, several of which have previously been implicated in abiotic stress signaling, hormonal regulation, and oxidative stress responses. The recurrent enrichment of these TF families supports the potential involvement of coordinated transcriptional control pathways during metal stress adaptation (**Table S9**). TMHMM analysis identified 32 genes encoding multi-pass transmembrane proteins, including transporters associated with Zn, Fe, Cu, phosphate, nitrate, sodium, and potassium transport (**Table S10**). Prominent transporter-associated CGs included *OsZIP2*, *OsZIP8*, *OsZIP10*, *OsCOPT7*, *OsFPN1*, *OsPEZ1*, *OsHKT4*, and *OsNRT2.3*, highlighting substantial enrichment of ion transport and membrane homeostasis functions within prioritized M-QTL regions. Several transporter-associated loci co-localized with genes involved in hormonal regulation, ROS buffering, and stress-responsive signaling, supporting potential coordination between transport-mediated detoxification and adaptive stress signaling processes. Subcellular localization analysis further revealed broad compartmentalization of prioritized proteins across the plasma membrane, vacuole, cytoplasm, endoplasmic reticulum, extracellular space, and nucleus (Table S11). Transport-associated proteins such as *OsZIP10*, *OsPIP2;5*, and *OsUMAMIT15* were predominantly localized to the plasma membrane, whereas *OsZIP2* and *OsZIP8* additionally exhibited predicted vacuolar or lysosomal localization, supporting possible roles in intracellular metal sequestration. Glutathione S-transferases and hormone-associated enzymes were primarily cytoplasmic, while several cytochrome P450 proteins localized to the endoplasmic reticulum. Extracellular localization was predicted for multiple lipid transfer proteins, peroxidases, and cell wall-associated proteins, including *OsLTPd3*, *OsPRX70*, *OsPRX77*, and *OsXTH21*, potentially linking these proteins to ROS scavenging and structural remodeling processes. Collectively, the predicted subcellular distribution patterns support compartment-specific coordination of transport, detoxification, signaling, and structural adaptation pathways during acidic soil-associated metal stress responses.

#### 3.4.5. Integrative genomics-led prioritization of M-QTLs for molecular breeding

The 98 prioritized candidate genes (CGs) showed uneven distribution across recurrent M-QTL intervals, indicating locus-specific enrichment of distinct stress-responsive functional modules (Fig. 5; Table S6). M-QTL 9.5 contained the highest density of prioritized CGs, followed by M-QTLs 10.9, 6.6, 10.7, and 11.1. Functional categorization of these regions revealed recurrent enrichment of pathways related to redox regulation, membrane transport, hormonal signaling, detoxification, and cellular homeostasis.

**Fig. 5.**
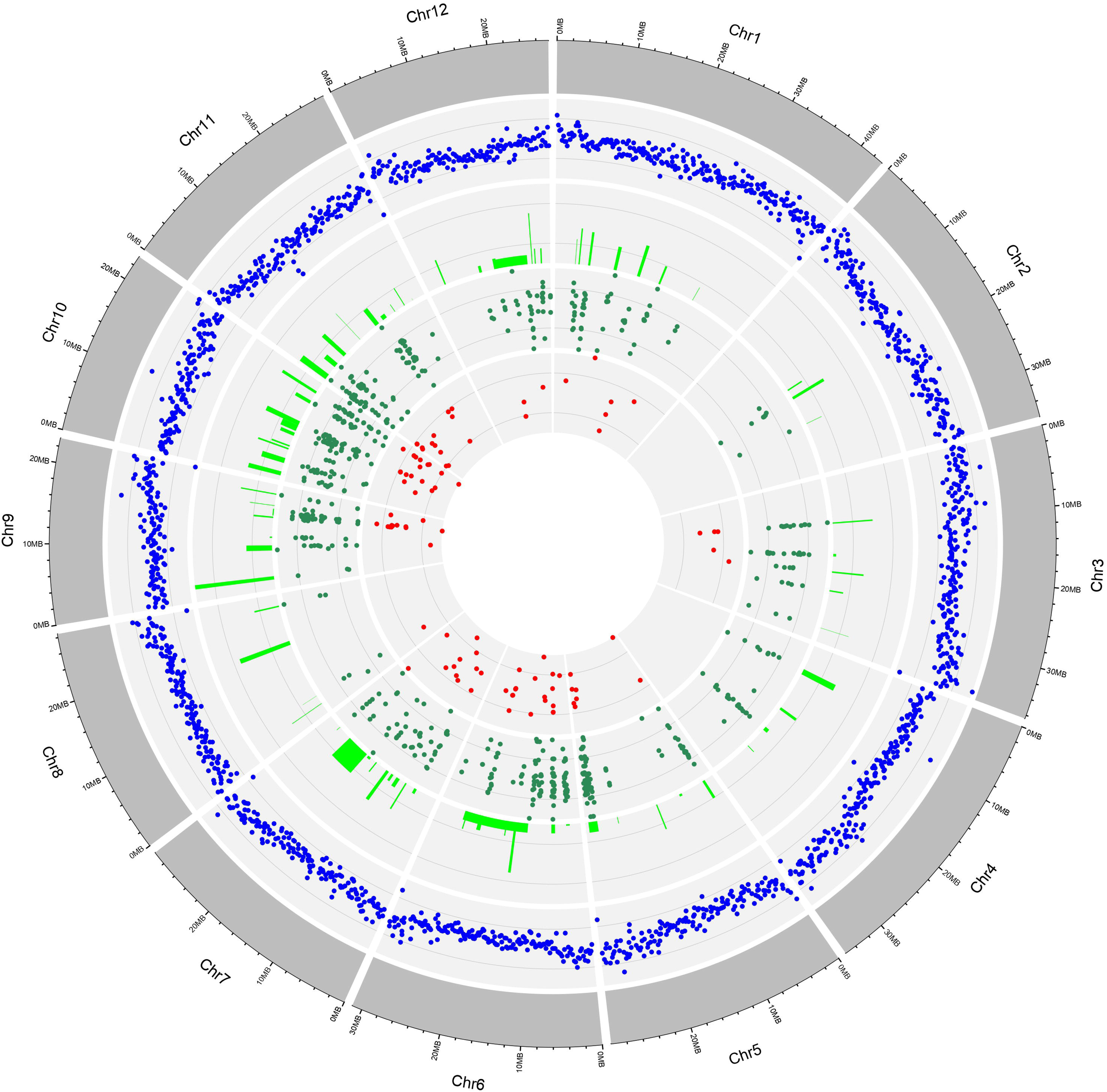
Distribution patterns of gene density, QTLs, M-QTLs, M-QTL-spanning genes, and candidate genes (CGs) identified on the rice genome. The outermost circle illustrates chromosome positions on the rice genome in Mb. The second circle, highlighted in blue, represents gene density across the rice genome. The third and fourth inner circles depict the number of initial AlCdMn toxicity tolerance QTLs and the identified M-QTLs genes, respectively. The fifth circle illustrates the underlying 98 prioritized candidate genes of the M-QTLs.

Among the prioritized regions, M-QTL 9.5 represented a prominent integrative hotspot containing transcriptional regulators (*OsbHLH120* and *OsERF102*), transport-associated genes (*OsCOPT7*, *OsSTP14*, and *OsPT19*), stress-responsive hub genes (*OsIDI4* and *OsUbL40-1*), and ethylene biosynthesis-associated genes (*OsACO1* and *OsACO2*). The co-occurrence of transporters, signaling-associated genes, and transcriptional regulators within this interval supports the potential involvement of coordinated transport and stress-signaling processes during metal stress adaptation.

M-QTL 10.9 was strongly enriched for glutathione S-transferase family genes, including *OsGSTU10*, *OsGSTU23*, *OsGSTU26*, *OsGSTU48*, and *OsGSTU50*, together with *OsHIPP25* and cysteine-rich stress-associated proteins, highlighting substantial representation of redox buffering and detoxification-associated pathways. Similarly, M-QTL 6.6 contained the metal transporter *OsZIP10* together with gibberellin biosynthesis-associated genes (*OsKO4* and *OsKO5*), potentially linking ion homeostasis with stress-responsive growth regulation. Additional recurrent M-QTLs contained genes associated with aquaporin-mediated transport, secondary metabolism, transcriptional regulation, and membrane-associated signaling functions.

Collectively, the prioritized M-QTLs represent recurrent genomic regions enriched for coordinated stress-responsive pathways involved in transport, detoxification, signaling, and cellular homeostasis. These loci provide biologically prioritized targets for downstream functional validation, marker development, and genomics-assisted breeding strategies for acidic soil-associated metal stress tolerance in rice.

### 3.5. Stress-dependent physiological and molecular responses of contrasting rice genotypes under individual and combined metal toxicities

#### 3.5.1. Differential root system and biomass adaptations

Root- and biomass-associated traits exhibited significant genotype, treatment, and genotype × treatment effects (p < 0.01), with root length (RL) showing the strongest interaction sensitivity (p < 0.001; Fig. S3–S5). Under control conditions, RL remained comparable between the two genotypes. However, under metal stress conditions, Sahasarang generally maintained higher RL than IR64, particularly under Mn and Al+Mn treatments, whereas IR64 showed pronounced root growth inhibition under several combined metal exposures, especially Al+Cd.

Similar genotype-dependent trends were observed for root surface area (SA), root volume (RV), root fresh weight (FW_R), and root dry weight (DW_R), although the magnitude of stress responses varied among treatment combinations. Overall, Sahasarang maintained comparatively greater root structural integrity under Al-containing treatments, whereas IR64 exhibited stronger reductions in root-associated traits under multiple combined stress conditions. Shoot biomass traits were comparatively less affected than root-associated parameters; however, genotype-specific differences remained evident, particularly under Cd-containing treatments. Notably, Al+Mn exposure promoted comparatively greater root and shoot growth than the corresponding individual stresses in both genotypes, indicating potentially non-additive interactions between the two stress conditions (**Fig. S3–S5**).

Correlation analysis revealed strong positive associations among major root architectural traits, including RL, SA, FA, and FW_R (r > 0.80), together with a positive relationship between root and shoot dry biomass (r = 0.74), supporting coordinated maintenance of root system architecture under stress conditions. Principal component analysis further separated the two genotypes under combined metal treatments, with Sahasarang clustering closer to control treatments and IR64 displaying greater phenotypic dispersion (**Fig. S6**). These patterns support comparatively higher phenotypic stability of Sahasarang under combined metal stress, whereas IR64 exhibited stronger stress-associated divergence in root and biomass traits.

#### 3.5.2. Differential physiological and biochemical responses

Sahasarang and IR64 showed contrasting regulation of redox homeostasis and carbon partitioning under metal stress. In IR64, shoot TPC increased under Cd stress (≈55% above control) but declined sharply under combined stresses, particularly Al+Mn and Cd+Mn (>45% reduction), while root phenolics were strongly reduced under Al+Cd+Mn. Correspondingly, shoot FRAP values in IR64 declined under all treatments, with the steepest reduction under Al+Mn, whereas root FRAP remained near control levels or showed only modest increases under Al, Al+Cd, and Cd+Mn, suggesting comparatively weaker antioxidant buffering under combined stress conditions.

In contrast, Sahasarang exhibited comparatively stronger and more sustained antioxidant activation. Shoot TPC increased consistently under Cd and Mn stress (up to ≈75% above control), while root phenolics remained relatively stable across treatments. FRAP responses were markedly stronger than in IR64, with shoot FRAP increasing by up to ≈30% under Cd and Al+Cd, and root FRAP nearly doubling under Cd and remaining elevated under Mn and combined stresses, consistent with enhanced inducible redox buffering under several stress combinations.

Carbon partitioning responses further differentiated the genotypes. In IR64, shoot total soluble sugars (TSS) increased marginally under Cd (≈10%) but declined sharply under Al+Cd+Mn (≈33%), while root TSS responses were inconsistent. Starch levels in IR64 decreased across most treatments, particularly in roots under Al and Cd stress (>30% reduction). By contrast, Sahasarang showed strong and coordinated sugar accumulation, with shoot TSS increasing under Cd and Al+Cd (up to ≈60%) and root TSS more than doubling under Mn, remaining elevated across treatments. Starch reserves in Sahasarang were largely maintained or enhanced, with increased shoot starch under Al+Mn and sustained root starch under Al+Cd and Al+Mn, suggesting comparatively improved metabolic adjustment under several combined stress treatments. **(Fig. 7a-d**).

Correlation analysis supported these contrasting strategies. In Sahasarang, phenolics, antioxidant capacity, and soluble sugars were positively coupled in roots (phenol–FRAP r = 0.42; phenol–TSS r = 0.63; FRAP–TSS r = 0.78) and shoots (phenol–FRAP r = 0.57; FRAP–TSS r = 0.65), while starch showed inverse relationships with defense traits (r ≈ −0.64), consistent with stress-associated carbon reallocation. In IR64, coordination was weaker or inconsistent (e.g., shoot phenol–TSS r = 0.76 but TSS–starch r = −0.52), indicating less coordinated biochemical responses under combined stress.**(Fig. S7e).**

#### 3.5.3. Metal accumulation and translocation under single and combined metal stress

ICP-OES analysis revealed pronounced genotype-dependent differences in metal accumulation and partitioning that became more evident under combined metal exposure (Fig. S8). Under single-metal treatments, IR64 accumulated comparatively high levels of Mn in roots (2502.50 µg g^-1^) and substantial Cd under Cd stress, whereas Sahasarang generally maintained lower Al and Cd accumulation while preserving Mn uptake capacity. These patterns indicate intrinsic differences in metal uptake and internal partitioning between the two genotypes.

Combined metal stress further accentuated these contrasts. In IR64, root Al and Cd accumulation increased markedly under Al+Cd treatment (Al: 2606.25 µg g^-1^; Cd: 583.75 µg g^-1^), while Cd accumulation increased further under Cd+Mn exposure (1192.17 µg g^-1^). Elevated root accumulation was accompanied by increased shoot translocation under Cd+Mn, particularly for Cd (664.00 µg g^-1^) and Mn (2850.00 µg g^-1^), indicating reduced restriction of metal movement from roots to shoots under combined stress conditions. In contrast, Sahasarang maintained comparatively lower shoot metal accumulation across combined treatments and retained a substantial fraction of Cd within roots under Al+Cd+Mn stress (856.39 µg g^-1^), while limiting shoot Al (158.00 µg g^-1^ and Mn (1528.00 µg g^-1^) accumulation.

Root-to-shoot translocation ratios (RSRs) further differentiated the genotypes. IR64 consistently exhibited higher Cd and Mn translocation under Cd, Cd+Mn, and Al+Cd+Mn treatments, whereas Sahasarang maintained lower RSRs across both single and combined stresses, indicating greater restriction of root-to-shoot metal translocation. Metal interaction analysis provided additional insight into these accumulation patterns. In IR64, Al uptake was strongly synergistic under Al+Cd (observed: 2606.25 vs. expected: 841.45 µg g^-1^) but antagonized under Al+Cd+Mn. Cd uptake was synergistically enhanced under Cd+Mn but suppressed by Al in Al-containing combinations. Mn uptake was strongly antagonized by both Al and Cd, with maximal suppression under triple stress (root: −1384.0 µg g^-1^; shoot: −2816.0 µg g^-1^ relative to expected). In contrast, Sahasarang maintained stable Al and Mn uptake under Al+Mn, restricted Al translocation under Al+Cd+Mn, and showed relatively stable Mn mobility across stress combinations.

Collectively, the ionomic profiles indicate that combined metal exposure amplified genotype-specific differences in metal partitioning behavior. IR64 exhibited greater accumulation and shoot translocation of toxic metals under several combined stress conditions, whereas Sahasarang maintained tighter control over root-to-shoot metal movement and lower shoot metal burden. The contrasting interaction patterns among Al, Cd, and Mn further highlight the interconnected nature of metal homeostasis under acidic soil-associated stress conditions.

#### 3.5.4. Cell wall compositional remodeling under combined metal stress assessed by FTIR

FTIR analysis revealed pronounced genotype-dependent differences in cell wall-associated spectral responses, particularly under combined Al+Cd+Mn stress (Fig. 6). In Sahasarang roots, combined metal exposure increased the intensity of lignin-associated (∼1596 cm^-1^) and polysaccharide/pectin-associated (1041–1086 cm^-1^) bands, accompanied by a slight red shift in the lignin peak (∼1596 to ∼1594 cm^-1^). These changes are consistent with altered cell wall composition involving lignin- and polysaccharide-associated spectral features. The O–H stretching region (3380–3405 cm^-1^) remained comparatively stable under combined stress, indicating maintenance of hydration-associated wall features. Similar but less pronounced spectral changes were observed under dual-metal treatments, whereas single-metal stresses induced comparatively minor modifications.

**Fig 6.**
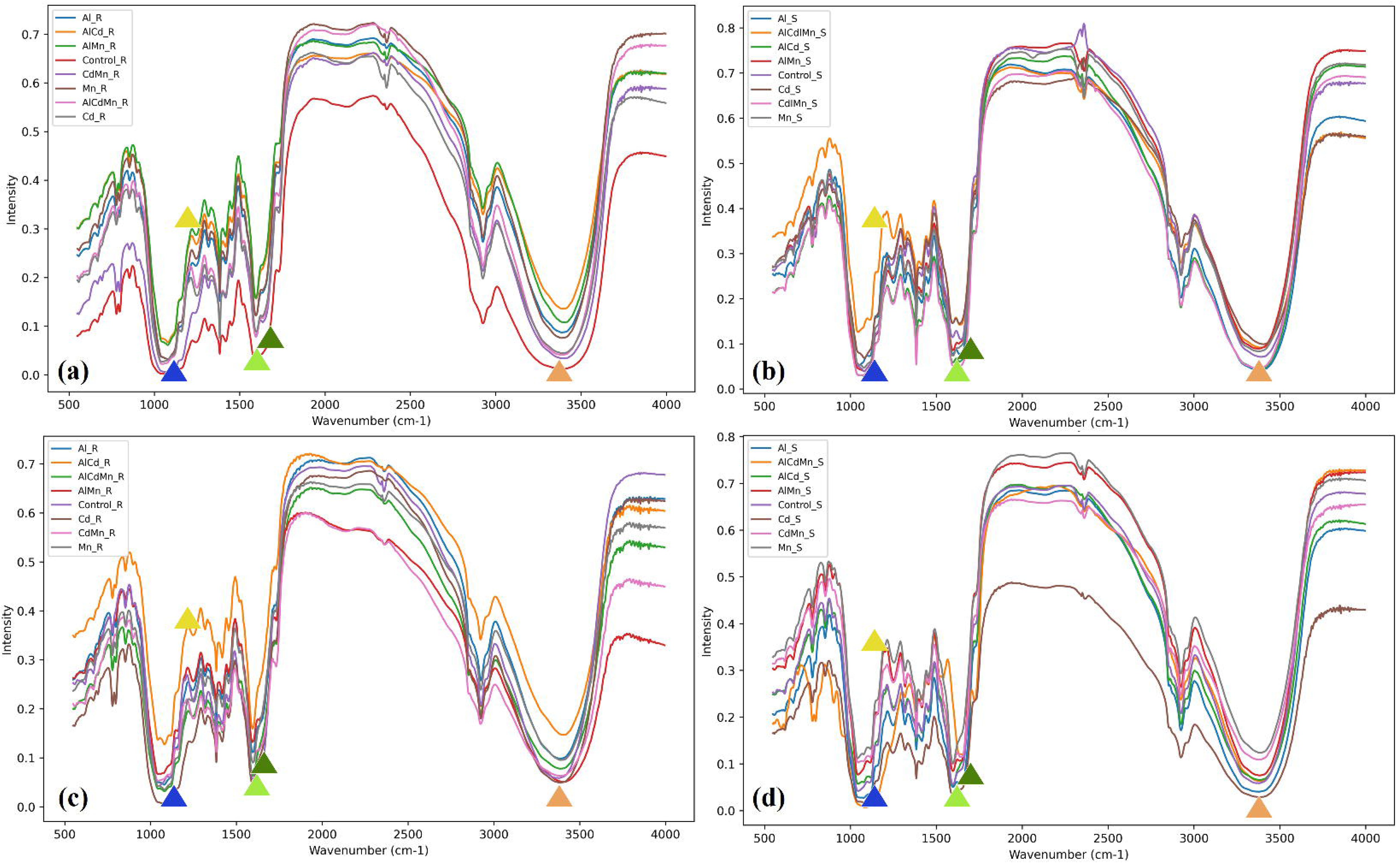
FTIR spectral profiles (4000–400 cm^-1^) of root and shoot tissues of rice genotypes IR64 and Shahsarang exposed to Al, Cd, Mn, and their combinations, showing key spectral changes in the O–H stretching region (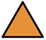∼3400 cm^-1^), the pectin-associated region (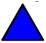∼1040–1086 cm^-1^), and the lignin-associated region (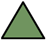∼1600 cm^-1^). (a) IR64 root, (b) IR64 shoot, (c) Shahsarang root, and (d) Shahsarang shoot. IR64 exhibited diminished O–H peaks, minimal pectin shifts, and weak or absent lignin signals under Cd and Al+Cd+Mn stress, whereas Shahsarang showed stable O–H bands, a consistent pectin shift (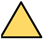∼1080 cm^-1^), and a distinct lignin peak (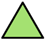∼1594 cm^-1^).

In contrast, IR64 roots exhibited substantial attenuation and broadening of lignin- and polysaccharide-associated bands under combined stress, together with a marked reduction in O–H stretching intensity. These spectral changes were most pronounced under Al+Cd+Mn treatment and indicate disruption of cell wall-associated structural organization under increasing stress complexity.

Shoot tissues showed patterns comparable to those observed in roots. Under combined stress, Sahasarang shoots retained strong lignin-associated (∼1594 cm^-1^) and pectin-associated (∼1080 cm^-1^) signals together with relatively stable O–H stretching profiles. By contrast, IR64 shoots displayed weak lignin-associated signals, limited polysaccharide-associated remodeling, and reduced O–H stretching intensity. Overall, the FTIR profiles indicate that Sahasarang maintained a more coordinated cell wall remodeling response under combined metal stress, whereas IR64 exhibited progressive disruption of cell wall-associated spectral features as stress complexity increased.

### 3.6. RT-qPCR-based expression profiling of prioritized candidate genes under individual and combined metal stresses

RT-qPCR analysis was conducted to profile prioritized CGs in roots and shoots of the acid-tolerant genotype Sahasarang and the sensitive genotype IR64 under individual (Al, Cd, Mn) and combined (Al+Cd, Al+Mn, Cd+Mn, Al+Cd+Mn) metal stresses (**Fig. 7; Fig. S9**). Based on their genotype-dependent expression behavior under combined stress and their association with M-QTL intervals enriched for stress-responsive functions, ten genes were retained for the main text *(OsACO1, OsACO4, OsKO4, OsJAZ13, OsZIP10, OsCOPT7, OsNrat1, OsbZIP50, OsbHLH120, OsGSTU10*). Additional responsive genes showing limited genotype discrimination (*OsACO2, OsDXS2, OsPIOX, OsLDOX2, OsUMAMIT15, OsLTP*) are presented in the Supplementary Information.

**Fig 7.**
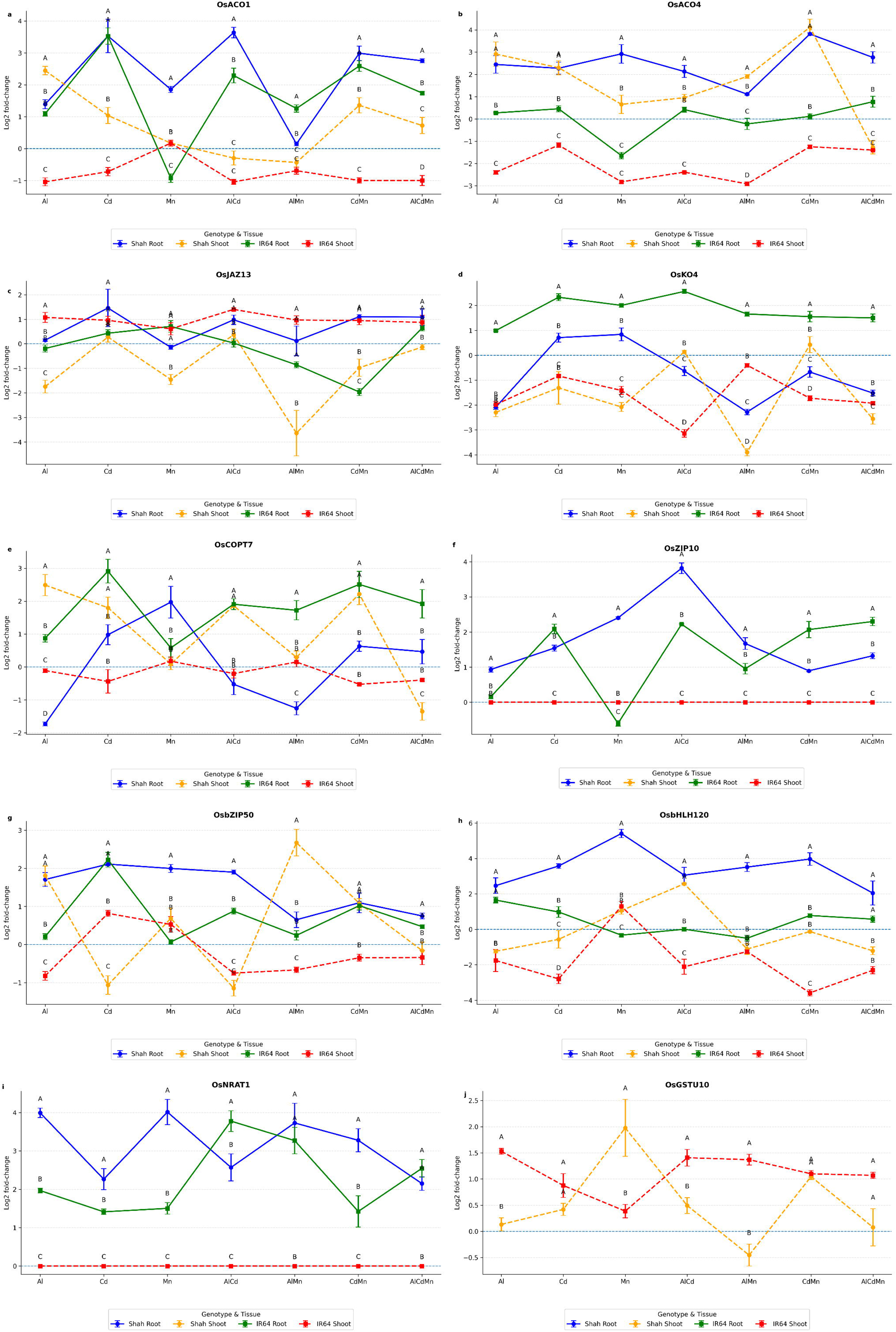
Expression profiles of candidate genes (CGs; A–J) in roots and shoots of tolerant (Shahsarang) and sensitive (IR64) rice genotypes under individual and combined metal stresses. Bar plots show log2 fold changes in transcript levels of *OsACO1, OsACO4, OsKO4, OsJAZ13, OsZIP10, OsCOPT7, OsNrat1, OsbZIP50, OsbHLH120, OsGSTU10*, as determined by RT-qPCR. Plants were subjected to Al, Cd, Mn, and their combinations. Expression was analyzed in both root and shoot tissues of each genotype. Bars represent the mean ± SE of three biological replicates. The differential expression patterns highlight genotype- and tissue-specific responses to metal toxicity, indicating coordinated transcriptional regulation underlying stress adaptation.

**Fig 8.**
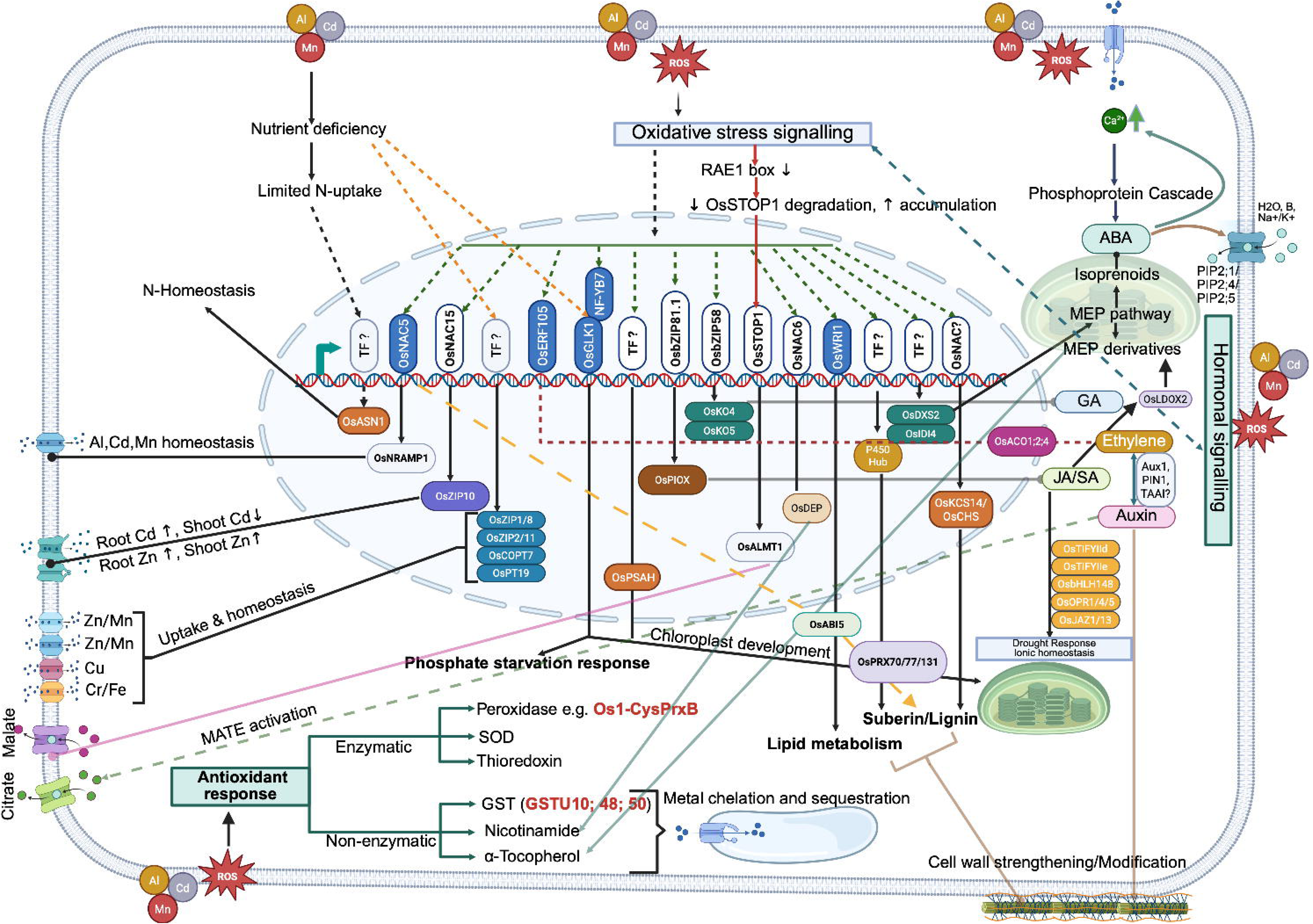
Rice metal stress response network emphasizing CGs identified in the current study. The colored text and boxes represented are among the high-confidence CGs.

Hormone-related genes showed strong genotype- and tissue-dependent regulation under combined metal stress. In roots, *OsACO1* was more strongly induced in Sahasarang than in IR64 under Cd-containing treatments and combinations (Cd, Al+Cd, Cd+Mn, Al+Cd+Mn), whereas responses under Al+Mn were weak and Mn alone induced moderate expression in both genotypes. *OsACO4* displayed clearer genotype dependent expression differences under combined stresses, while *OsKO*4 showed opposite regulation between genotypes: Sahasarang consistently downregulated *OsKO4* under all Al-containing treatments, including combinations, whereas IR64 showed induction across most single and combined stresses. In shoots, *OsJAZ13* was reduced in Sahasarang under combined stresses (notably Al+Mn and Cd+Mn), while IR64 showed consistent induction across treatments.

Metal transporter genes exhibited predominantly root-centered regulation. *OsZIP10* showed consistent upregulation in Sahasarang roots across all single and combined treatments, whereas IR64 displayed variable induction, largely confined to Cd stress and reduced under Mn-containing combinations. *OsCOPT7* showed genotype-dependent regulation in roots, being downregulated under Al-containing treatments and induced under Cd- or Mn-containing treatments in Sahasarang, but broadly upregulated across treatments in IR64. The benchmark Al transporter *OsNrat1* showed strong and consistent root-specific induction in both genotypes under all Al-containing treatments, with minimal shoot expression, confirming its conserved role in Al detoxification.

Among transcription factors, *OsbHLH12*0 exhibited higher expression in Sahasarang roots under combined metal stresses, whereas IR64 showed weaker induction. *OsbZIP50* showed moderate, treatment-dependent induction, with higher root expression in Sahasarang under Cd- and Mn-containing combinations and low, variable shoot expression in both genotypes. Metabolic and detoxification-related genes showed tissue bias: *OsGSTU10* was predominantly induced in shoots, with higher expression in IR64 across treatments, while *OsDXS2* showed strong shoot induction in IR64 but increased root expression in Sahasarang under Mn and Cd+Mn stress.

Overall, CGs derived from multimetal stress M-QTLs showed clearer differences in expression magnitude, tissue specificity, and genotype dependence under combined metal stress than under single-metal treatments. These patterns indicate that stress complexity unmasks regulatory divergence between Sahasarang and IR64, consistent with their contrasting stress response phenotypes. Although the observed transcriptional divergence provides supportive evidence for involvement of these CGs in metal stress responses, expression-based analyses alone do not establish causal function. Therefore, definitive validation of the prioritized candidates will require targeted reverse-genetic and functional characterization approaches.

## 4. Discussion

### 4.1. Meta-QTL analysis

Soil acidity frequently exposes rice to concurrent Al, Cd, and Mn toxicities, particularly in the upland systems (Haefele *et al*., 2014; Zhao *et al*., 2014; Bailey *et al*., 1995; Jung *et al*., 2008; Sarwar *et al*., 2010; Gao *et al*., 2022). This underscores the importance of understanding both shared and stress-specific adaptive responses under acidic soil conditions where multiple toxic metals frequently co-occur. Although numerous QTLs and MTAs for Al, Cd, and Mn stress response have been reported, their deployment in breeding has been limited (Awasthi *et al*., 2017; Tao *et al*., 2018; Rasheed *et al*., 2020). Notably, 681 QTLs/MTAs have been reported altogether for these metals through the 37 QTL and 16 association mapping approaches as mentioned in the introduction. The availability of these large-scale datasets provides an opportunity to identify robust genomic regions associated with rice responses to acidic soil-associated metal stresses across independent studies.

Using these datasets, we conducted an M-QTL analysis and successfully projected 493 (72.39%) QTLs and MTAs associated with Al, Cd, and Mn stress responses onto a reference linkage map spanning 1837.97 cM, reflecting strong marker correspondence and overall dataset consistency (Goffinet & Gerber, 2000; Veyrieras et al., 2007). These loci were consolidated into 79 M-QTLs, supporting a complex and polygenic genetic architecture underlying rice responses to metal toxicity.. Chromosomes 7 and 10 emerged as major genomic hotspots, followed by chromosome 11, consistent with their reported enrichment for abiotic stress–related loci in cereals (Kumar *et al*., 2015; Khahani *et al*., 2021) **(Table S3)**.

Nearly one-third of the loci (23/79) contained ≥5 overlapping QTLs, indicating their recurrent detection across diverse mapping studies (Kumari *et al*., 2023, 2024). Notably, ten M-QTLs were associated with traits linked to all three metals and explained 14.8–38.25% of the phenotypic variance across integrated datasets. These regions represent priority genomic regions associated with interconnected transport, detoxification, and stress-responsive regulatory pathways under acidic soil-associated metal stress (Courtois *et al*., 2009; Khahani *et al*., 2021). Likewise, M-QTLs 7.2 and 7.4 showed the highest QTL density and PVE, and may represent regions containing multiple stress-associated loci contributing to the observed clustering of QTLs. The recurrent clustering of QTLs with comparatively high PVE values across independent studies indicates that these regions likely represent stable components of multimetal stress adaptation across diverse genetic backgrounds.

One of the major outcomes of the present analysis was the substantial improvement in mapping resolution, with the mean confidence interval reduced from 6.90 cM to 1.85 cM, thereby improving CG resolution within stress-associated loci.. Several identified M-QTLs, including M-QTL 5.5 and M-QTL 6.3, harbored multiple CGs associated with metal transport, detoxification, and stress-responsive signaling pathways. Several M-QTLs therefore represent high-priority regions for dissecting the genetic basis of ion transport regulation, detoxification, and stress-responsive signaling under combined metal stress. Importantly, the phenotypic traits integrated in the present meta-analysis represented distinct but biologically interconnected manifestations of metal stress adaptation. Al-associated studies predominantly evaluated root growth inhibition traits, whereas many Cd- and Mn-associated studies focused on tissue accumulation, partitioning, and translocation parameters that are widely used as indicators of metal toxicity tolerance and detoxification efficiency in rice. Thus, overlapping M-QTLs likely reflect broader stress-associated adaptive processes operating across distinct physiological contexts.

### 4.2. Prioritization of CGs

Integration of refined M-QTL intervals with a multi-criteria prioritization framework enabled the identification of high-confidence CGs associated with Al, Cd, and Mn stress responses. Rather than relying solely on differential expression, genes were prioritized using a composite score integrating recurrence across independent genetic studies, expression behaviour across stress datasets, functional annotation, and M-QTL support. Combined analysis of positional information, functional annotation, and meta-transcriptomic evidence from publicly available RNA-seq datasets yielded 98 high-confidence CGs supported across independent studies.

Promoter analysis of these 98 CGs revealed extensive enrichment of stress- and hormone-responsive cis-regulatory elements. A high proportion of CGs harboured MYB or MYB-related binding sites (86.7%), ABREs (80.6%), and STRE/G-box elements (78.6%), while DRE (53.1%) and AUXRR (13.3%) were also represented. The observed cis-element composition is consistent with ABA-, MYB-, and stress-responsive regulatory pathways frequently associated with oxidative and metal stress responses (Guiltinan *et al*., 1990; Gallego *et al*., 2012; Liu *et al*., 2016; Shukla *et al*., 2015). In addition, several transcription factor families, including NAC, AP2/ERF, bHLH, and G2-like proteins, were represented among the prioritized candidates, highlighting extensive representation of transcriptional regulators among prioritized candidates (Dubouzet *et al*., 2003; Narusaka *et al*., 2003; Arenhart *et al*., 2016; Tang *et al*., 2023; Ofoe *et al*., 2023; Anwar *et al*., 2024). Transport-associated candidates represented a major component of the prioritized CG set, including *OsZIP2/8/10*, *OsCOPT7*, *OsFPN1*, *OsPT19*, *OsPEZ1*, *OsHAK24*, and *OsNRT2.3*, all previously associated with metal or inorganic ion homeostasis under stress conditions (Hanikenne *et al*., 2008; Wong & Cobbett, 2009; Drew *et al*., 2021). Conversely, several canonical metal transporters, including OsNramp1 and OsNramp5, were not retained, likely reflecting limitations related to QTL resolution, conditional regulation, or spatiotemporal specificity of expression rather than absence of biological significance (Ishimaru *et al*., 2012; Sasaki *et al*., 2012; Chang *et al*., 2020).

Subcellular localization predictions further indicated compartment-specific functional organization among the prioritized CGs, including vacuolar metal sequestration-associated transporters (*OsZIP2* and *OsZIP8*), cytoplasmic detoxification- and hormone-associated proteins (OsGSTU10 and OsACO1), endoplasmic reticulum-localized cytochrome P450s, and nucleus-localized transcription factors such as *OsGLK1* and *OsNAC5*.. Notably, *OsACO4* exhibited predicted dual localization in the cytoplasm and nucleus, supporting potential functional complexity in ethylene-associated stress signaling. Protein–protein interaction analysis identified a highly interconnected network of hub genes associated with redox regulation, metabolic adjustment, and protein quality control under metal stress conditions. Central interaction nodes included redox-associated proteins such as 1-Cys peroxiredoxin B, metabolic enzymes linked to nitrogen, methionine, lipid, and isoprenoid metabolism (e.g., OsIDI4, CYP86B1, and DXS2/3), and components associated with ubiquitin-mediated protein turnover (e.g., UbL40-1). These interaction modules are consistent with involvement of redox regulation, metabolic plasticity, and proteostasis-associated pathways in rice responses to acidic soil-associated metal stresses (Compagnon *et al*., 2009; Dansana *et al*., 2014; Kumar & Trivedi, 2018; Hu *et al*., 2020).

Overall, 4,702 non-redundant genes across 79 M-QTLs were refined to 98 high-confidence candidates distributed across 39 metal stress–responsive M-QTLs, with major hotspots at M-QTL 9.5, 10.9, 6.6, and 11.1. Together, this multi-parametric prioritization framework defines a biologically supported CGs set that provides a foundation for future functional characterization and genomic-assisted breeding efforts targeting acidic soil-associated metal stress adaptation. (**Table S3**; **Fig. 5**). Although the prioritized CGs were supported by convergent positional, transcriptional, and functional evidence, reverse-genetic and physiological validation will be required to establish their precise mechanistic roles in stress adaptation.

### 4.3. Comparative evaluation of Sahasarang and IR64 under individual and combined Al, Cd, and Mn stress

Physiological evaluation of Sahasarang and IR64 was undertaken to characterize genotype-dependent responses to individual and combined metal stress and to provide biological context for the stress-responsive pathways represented within the prioritized candidate gene set. Under several combined metal treatments, Sahasarang maintained greater root system integrity, including higher root volume, surface area, and biomass relative to IR64, particularly under Al-containing stress combinations. In contrast, IR64 exhibited stronger reductions in root-associated traits under several combined stress conditions. Maintenance of root volume, surface area, and dry biomass in Sahasarang is consistent with enhanced structural stability under metal stress conditions, traits frequently associated with adaptation to acidic and metal-affected soils (Wang *et al*., 2023a; Tang *et al*., 2024; Qin *et al*., 2024). The relatively stable multivariate phenotypic profile of Sahasarang across combined stress treatments further supports substantial genotype-dependent divergence in multimetal stress adaptation. Physiological and biochemical analyses further highlighted contrasting genotype-dependent stress responses. Sahasarang exhibited comparatively stronger accumulation of soluble sugars, phenolics, and antioxidant-associated responses under several combined stress treatments, consistent with stronger metabolic buffering and antioxidant-associated stress acclimation. In contrast, IR64 showed more variable antioxidant responses across treatments, with pronounced reductions under several combined stress conditions, indicating less stable antioxidant and metabolic regulation under increasing stress complexity. Cell wall remodeling emerged as an important differentiating response between the two genotypes. Cell wall-associated immobilization and apoplastic sequestration represent major components of plant metal detoxification responses, with lignin and pectin playing central roles in metal binding and immobilization (Bhuiyan *et al*., 2007; Moura *et al*., 2010; Amos and Mohnen, 2019; Riaz *et al*., 2021; Wu *et al*., 2021; Lin *et al*., 2022). FTIR signatures in Sahasarang under Al+Cd+Mn stress indicated stronger lignification and polysaccharide-related spectral responses together with preservation of apoplastic hydration, consistent with altered cell wall properties and modified metal interaction dynamics under combined stress (Carrasco-Gil *et al*., 2013; Kopittke *et al*., 2015; Xiong *et al*., 2009). In contrast, attenuation of lignocellulosic and O–H-associated bands in IR64 indicated greater disruption of cell wall-associated spectral features under combined stress, potentially exacerbated by oxidative damage and lipid peroxidation (Rizwan e*t al*., 2016).

Elemental profiling further supported these observations, as IR64 accumulated comparatively higher concentrations of Al, Cd, and Mn in roots and shoots and exhibited elevated root-to-shoot translocation ratios relative to Sahasarang under several stress combinations. Both genotypes showed reduced Mn uptake in the presence of Al or Cd, consistent with antagonistic metal–metal interactions, whereas Sahasarang generally maintained more stable metal partitioning patterns across treatments. These findings align with classical reports of Al–Mn and Al–Cd antagonism (Rees and Sidrak, 1961; Clark, 1977; Blair and Taylor, 1997; Taylor *et al*., 1998; Yang *et al*., 2009; Shamsi *et al*., 2007), while highlighting strong genotype dependence in metal interaction responses. The partial alleviation of Al-and Cd-induced root inhibition by Mn is also consistent with earlier observations (Muhammad et al., 2016).

Compared with IR64, Sahasarang maintained more stable integration among root structure, antioxidant capacity, phenolic metabolism, and carbon reserve maintenance, consistent with contrasting stress-response architectures between the two genotypes. Collectively, these findings indicate that genotype responses to combined metal stress involve coordinated changes in root architecture, metabolic regulation, cell wall-associated responses, and metal partitioning, with Sahasarang generally exhibiting greater physiological stability than IR64 under several combined stress conditions. Importantly, genotype responses varied depending on stress composition, indicating that tolerance-associated responses to Al, Cd, and Mn are only partially overlapping. While Sahasarang exhibited comparatively stronger performance under several Al-containing and combined stress treatments, responses under Cd-associated treatments were more variable, suggesting that adaptation to acidic soils does not necessarily confer equivalent protection against all metal toxicities. These observations further support substantial genotype- and metal-dependent divergence in stress responses under combined metal exposure.

### 4.4. RT-qPCR-based validation of CGs for Al, Cd, and Mn toxicity tolerance

Expression profiling of Sahasarang and IR64 under individual and combined metal treatments revealed strong genotype-, tissue-, and stress-combination–dependent transcriptional divergence. Hormone-related CGs exhibited strong genotype-, tissue-, and stress-combination–dependent regulation, particularly under combined metal stress conditions. *OsACO1* (Cd-containing treatments and combinations) and *OsACO4* (across treatments) exhibited stronger and more consistent induction in Sahasarang, consistent with differential ethylene-associated transcriptional regulation between genotypes under combined stress. Ethylene-associated pathways have previously been implicated in regulation of lignification, apoplastic modifications, and stress-responsive signaling under metal toxicity conditions (Vassilev *et al*., 2004; Rodríguez-Serrano *et al*., 2006; Rzewuski and Sauter, 2008; Steffens, 2014; Wang *et al*., 2015). In contrast, IR64 displayed lower and more variable *OsACOs* expression across treatments, suggesting contrasting ethylene-associated transcriptional responses between the two genotypes. The gibberellin biosynthesis gene *OsKO4* showed consistent repression across treatments, with stronger and more uniform downregulation in Sahasarang consistent with previous findings (Lu *et al*., 2024). The coordinated repression of *OsKO4* alongside enhanced ethylene-associated signaling in Sahasarang is consistent with hormone-mediated growth–defense reprogramming, in agreement with reported DELLA- and EIN-dependent ethylene–GA crosstalk (Achard *et al*., 2003; Weiss and Ori, 2007). These responses may be associated with the comparatively improved plant architecture observed under Al+Mn treatment, although the underlying molecular mechanisms remain unresolved.

Likewise, *OsJAZ13*, a repressor of JA signaling (Feng *et al*., 2020), was reduced in Sahasarang shoots compared to IR64 under several combined metal treatments, potentially reflecting altered jasmonate-associated regulatory responses under combined stress. Notably, JA- and lipid-associated stress-responsive genes such as *OsPIOX* showed pronounced shoot induction in IR64, consistent with broader stress-responsive activation patterns relative to Sahasarang,, and are therefore presented as supplementary evidence.

Among transcription factors, *OsbHLH120* showed strong root induction in Sahasarang under combined metal stress, whereas IR64 responded weakly, extending its known roles in PEG-, salt-, and ABA-mediated stress responses to complex metal stress environments and suggesting possible involvement in root-associated stress responses under combined metal exposure (Li *et al*., 2015). *OsbZIP50* displayed moderate, treatment-dependent induction, with relatively higher expression in Sahasarang roots under Cd- and Mn-containing combinations. *OsbZIP50* activates *OsZIP10*, which is essential for maintaining Zn, Fe, and Cu homeostasis (Qing *et al*., 2024), and functions in the endoplasmic reticulum (ER) stress response in rice (Takahashi *et al*., 2012; Hayashi *et al*., 2012; Hasan *et al*., 2017), consistent with a possible association between ionic homeostasis and ER stress responses under metal toxicity.

Metal transporter CGs showed clear genotype-dependent regulation concentrated in roots, underscoring their role in metal sensing and detoxification. *OsZIP10*, a downstream target of the Cd/Zn-responsive TF OsNAC15 (Lilay *et al*., 2020; Kumar, 2021; Zhan *et al*., 2022), displayed stable induction across treatments in Sahasarang roots, consistent with differential regulation of micronutrient transport processes under combined stress, whereas IR64 exhibited more variable regulation, particularly under Mn-containing combinations. *OsCOPT7* exhibited contrasting regulation between genotypes, with repression in Sahasarang roots under Al-containing treatments but broad induction in IR64, indicating differential control of Cu homeostasis. OsNrat1, an Al-specific transporter mediating vacuolar sequestration of cytosolic Al (Xia *et al*., 2010), was strongly induced in Sahasarang roots under Al stress and also upregulated under Cd and Mn treatments, indicating that components of Al-responsive transcriptional regulation may also respond under broader multimetal stress conditions. Collectively, these patterns highlight genotype-dependent transporter regulation under combined metal stress conditions.

Genes involved in detoxification and metabolic adjustment further differentiated genotype-specific stress-response patterns. *OsGSTU10*, associated with oxidative stress detoxification and previously implicated in abiotic stress responses including cold and herbicide exposure (Jain *et al*., 2010; Martini *et al*., 2022), was strongly upregulated in shoots of both genotypes under most metal stresses, except under Al and Al+Mn treatments in Shahsarang, where expression was slightly reduced, indicating that antioxidant-associated transcriptional responses differed depending on stress composition and genotype. *OsDXS2*, encoding the first committed enzyme of the methylerythritol 4-phosphate (MEP) pathway, exhibited pronounced genotype- and tissue-specific regulation, with consistently higher shoot expression in IR64 across treatments, indicative of generalized stress activation, whereas Shahsarang showed lower shoot expression and moderate, treatment-dependent root induction, indicating differential tissue-specific expression dynamics between genotypes.

One of the primary objectives of this study was to test whether CGs derived from multi-metal meta-QTLs exhibit genotype-dependent transcriptional divergence under combined metal stress. Across the CG set, dual and triple metal treatments resolved markedly clearer differences in expression magnitude, tissue specificity, and genotype dependence between Sahasarang and IR64 than single-metal exposures. The heterogeneous CG expression reflects context-dependent, genotype-specific regulation under combined metal stress rather than inconsistent candidate behaviour.

At the transcriptional level, Sahasarang displayed comparatively selective and predominantly root-centered regulation of CGs involved in hormone signaling, metal transport, transcriptional control, and metabolic adjustment, whereas IR64 showed broader and less tissue-specific transcriptional activation, particularly in shoots, consistent with its contrasting physiological response profile under combined stress. These observations further support the idea that tissue-specific regulatory organization, rather than expression magnitude alone, contributes to contrasting genotype responses under combined metal stress.

These findings align with extensive evidence that transcriptional responses to concurrent stresses are not predictable from single-stress datasets, as combined stresses activate distinct regulatory networks rather than additive responses (Rasmussen *et al*., 2013; Pandey *et al*., 2015; Shaar-Moshe *et al*., 2017). Consistent with this paradigm, multi-metal treatments in the present study provided stronger genotype discrimination and clearer tissue-specific regulatory divergence among M-QTL CGs than individual metal stresses, highlighting the importance of combined-stress frameworks for resolving genotype-dependent regulatory responses not evident under individual metal treatments. (Rasmussen *et al*., 2013; Pandey *et al*., 2015). Although the observed transcriptional divergence strongly supports association of these CGs with metal stress responses, reverse-genetic and physiological validation will be necessary to establish their precise mechanistic roles under individual and combined metal stress conditions.

## Conclusion

This study establishes an integrative genomic framework for investigating rice responses to acidic soil-associated metal stresses through large-scale meta-QTL analysis. Integration of 681 reported QTLs and MTAs from 53 independent studies resolved 79 stable M-QTLs with substantially reduced confidence intervals, identifying key genomic regions associated with responses to aluminium, cadmium, and manganese stress. These findings support the complex and polygenic nature of rice adaptation to acidic and metal-stressed environments and provide a systems-level framework for investigating stress-adaptive responses under combined metal stress conditions.

Integration of refined M-QTL intervals with a multi-criteria CGs prioritization strategy enabled identification of high-confidence stress-associated genes enriched for metal transport, cell wall-associated responses, redox regulation, and hormone-related pathways. Physiological, biochemical, FTIR-based cell wall, ionomic, and transcriptional analyses of contrasting rice genotypes under individual and combined metal stresses provided biological support for these enriched pathways and revealed coordinated structural, metabolic, and molecular responses under complex stress conditions. Notably, genotype-dependent transcriptional divergence was substantially stronger under combined metal stress than under individual metal exposure, emphasizing the importance of multi-stress experimental frameworks for resolving complex stress-adaptive responses in rice. Although expression-based analyses provide associative rather than causal evidence, the prioritized M-QTLs and CGs identified here represent high-confidence targets for future allele mining, functional validation, and molecular breeding efforts aimed at improving adaptation to acidic soil-associated metal stress. Collectively, this work integrates population-scale meta-QTL analysis with physiological and molecular characterization of genotype-dependent stress responses, thereby linking genomic architecture with adaptive responses under combined metal stress conditions. Nevertheless, the integrated Al-, Cd-, and Mn-associated datasets represent distinct but biologically interconnected manifestations of metal stress adaptation. Accordingly, overlapping M-QTLs identified across these datasets are more appropriately interpreted as shared stress-associated genomic regions contributing to broader ion homeostasis, detoxification, and adaptive regulatory processes rather than evidence of identical tolerance mechanisms across all three metals.

## Authors’ contribution

All authors contributed to the study conception and design. Material preparation, data collection and analysis were performed by Sandeep Jaiswal, Anita Kumar and Kuldeep Kumar, Saurav Kumar, Santosh Kumar, Simardeep Kaur, Nitish Ranjan Prakash, Pankaj Baiswar, Alka Bharati, Manjeet Talukdar, Sanjay Behera. The first draft of the manuscript was written by Sandeep Jaiswal, Anita Kumar, Kuldeep Kumar, Binay K Singh and all authors commented on previous versions of the manuscript. All authors read and approved the final manuscript.

## Data Availability

The datasets generated during and/or analysed during the current study are available from the corresponding author on reasonable request.

## Ethics approval

Not applicable

## Consent to participate

Not applicable

## Consent to publish

Not applicable

## Acknowledgment

The authors gratefully acknowledge all collaborators and co-principal investigators of the institute-funded project “Genome-wide association study (GWAS) for Al toxicity tolerance in rice from the North Eastern Hill Region of India” (Project Code IXX17233) for their contributions. We thank Dr. Krishnappa Rangappa for access to the root phenotyping facility; Dr. Samir Das and Dr. A.A.P. Milton for laboratory support; Dr. Renu Pandey for assistance with ICP-OES analysis; and Dr. Bijen Singh for support with FTIR spectroscopy. We also appreciate the contributions of technical staff and colleagues involved in experimentation and data analysis, and thank the Director, ICAR Research Complex for NEH Region, Umiam, for providing the necessary facilities.

## Figure Legends

**Fig. S1.** Chromosome-wise distribution of QTLs for AlCdMn toxicity tolerance in rice used for M-QTL analysis (Blue). Red bars indicate the proportion of QTLs projected during the analysis while yellow bars indicate M-QTLs.

**Fig. S2.** Gene Ontology (GO) enrichment analysis results depicting the biological processes associated with 98 prioritized candidate genes.

**Fig. S3.** Representative phenotypes of rice seedlings (IR64, top row; Shahsarang, bottom row) under control and various single and combined metal stress treatments, as labeled. Differences in root and shoot growth highlight genotype-specific responses. Scale bar at left.

**Fig. S4.** Root and shoot trait responses of tolerant (Shahsarang) and sensitive (IR64) rice genotypes under control (C), individual (Al, Cd, Mn), and combined (Al+Cd, Cd+Mn, Al+Mn, Cd+Al+Mn) metal stress treatments. Traits include root length, surface area (SA), fork angle (FA), volume (Vol), average diameter (Avg D), fresh weight, dry weight, and shoot fresh and dry weights. Values represent mean ± SE (n = 3).

**Fig. S5.** Correlation of metal stress treatments (C, Cd, Al, Mn, CdAl, CdMn, AlMn, CdAlMn) with root and shoot traits in Shahsarang (left) and IR64 (right). Heatmaps show point-biserial correlation coefficients (r); red and blue indicate positive and negative correlations, respectively. Significance: **\***p < 0.05, **\*\***p < 0.01, **\*p < 0.001.**

**Fig. S6.** Principal component analysis (PCA) biplots of root system architecture and biomass traits in rice seedlings of Shahsarang (top row) and IR64 (bottom row) under control and individual or combined Al, Cd, and Mn stress treatments. Plots show trait loadings and sample distributions across PC1 vs PC2 (left) and PC1 vs PC3 (right). Traits include total root length (RL), root surface area (SA), number of forks (FA), root volume (RV), average root diameter (Avg_D), and fresh and dry weights of root (FW_R, DW_R) and shoot (FW_S, DW_S).

**Fig S7.** Biochemical responses in root and shoot tissues of rice genotypes IR64 and Shahsarang under single and combined metal stress treatments. Panels show (a) total soluble sugar (TSS), (b) starch, (c) phenol, and (d) FRAP (Ferric Reducing Antioxidant Power) content in roots and shoots of IR64 and Shahsarang across different treatments: control (C), aluminum (Al), cadmium (Cd), manganese (Mn), and their combinations (Al+Cd, Al+Mn, Cd+Mn, Al+Cd+Mn). Bars represent mean ± SD for each genotype–tissue group: IR64-Root (green), IR64-Shoot (orange), Shahsarang -Root (blue), and Shahsarang -Shoot (pink). Panel (**e)** Correlation heatmap showing pairwise Pearson correlations among biochemical traits [FRAP, phenol, starch, and total soluble sugars (TSS)] in root and shoot tissues of IR64 and Shahsarang. Color scale indicates correlation strength (–1.0 to +1.0).

**Fig. S8. a.** Accumulation of Al, Cd, and Mn in roots and shoots of tolerant (G4, Shahsarang) and sensitive (G2, IR64) rice genotypes under control (C), single (Al, Cd, Mn), and combined (Al+Cd, Cd+Mn, Al+Mn, Al+Cd+Mn) metal stress treatments. Values represent mean of three biological replicates. **b.** Root-to-shoot translocation ratios of Al, Cd, and Mn under single (Al, Cd, Mn) and combined (Al+Cd, Cd+Mn, Al+Mn, Al+Cd+Mn) metal stress treatments in Shahsarang and IR64. Values represent means of three biological replicates.

**Fig. S9.** Expression profiles of candidate genes (CGs; K–P) in roots and shoots of tolerant (Shahsarang) and sensitive (IR64) rice genotypes under individual and combined metal stresses. Bar plots show log2 fold changes in transcript levels of *OsACO2, OsLDOX, OsPIOX, OsUMAMIT15, OsDXS, and OsLTP*, as determined by RT-qPCR.

## Table Legends

**Table S1.** List of PCR primers used for the RT-qPCR-based expression analysis of selected high confidence candidate genes in Shahsarang and IR64.

**Table S2.** Summary of QTL mapping studies on Al, Cd, and Mn toxicity tolerance in rice used for meta-analysis.

**Table S3.** List of 79 M-QTLs shortlisted based on the number of overlapping QTLs.

**Table S4.** Details of M-QTLs identified for toxicity tolerance to all three metals (Al, Cd, and Mn).

**Table S5.** Summary of 4,702 non-redundant genes identified within the 79 M-QTL regions.

**Table S6.** List of prioritized candidate genes derived from M-QTL

**Table S7.** Top 10 STRING network ranked by MCC method.

**Table S8.** CG wise cis-element matrix of prioritized CGs.

**Table S9.** List of transcription factors predicted from the CGs.

**Table S10.** List of 32 unique genes encoding multi-pass transmembrane proteins (≥3 TM helices).

**Table S11.** Subcellular localization predictions of prioritized CGs

